# PAM binding ensures orientational integration during Cas4-Cas1-Cas2 mediated CRISPR adaptation

**DOI:** 10.1101/2022.05.30.494039

**Authors:** Yukti Dhingra, Shravanti K. Suresh, Puneet Juneja, Dipali G. Sashital

## Abstract

Adaptation in CRISPR-Cas systems immunizes bacteria and archaea against mobile genetic elements. In many DNA-targeting systems, the Cas4-Cas1-Cas2 complex is required for selection and processing of DNA segments containing PAM sequences, prior to integration of these “prespacer” substrates as spacers in the CRISPR array. We determined cryo-EM structures of the Cas4-Cas1-Cas2 adaptation complex from the type I-C system that encodes standalone Cas1 and Cas4 proteins. The structures reveal how Cas4 specifically reads out bases within the PAM sequence and how interactions with both Cas1 and Cas2 activate Cas4 endonuclease activity. The Cas4-PAM interaction ensures tight binding between the adaptation complex and the prespacer, significantly enhancing integration of the non-PAM end into the CRISPR array and ensuring correct spacer orientation. Corroborated with our biochemical results, Cas4-Cas1-Cas2 structures with substrates representing various stages of CRISPR adaptation reveal a temporally resolved mechanism for maturation and integration of functional spacers into the CRISPR array.

## Introduction

CRISPR (clustered regularly interspersed palindromic repeats) arrays and Cas (CRISPR associated) proteins constitute adaptive and heritable immune systems that rely on molecular recordings of pathogen invasion, enabling rapid and specific response to future infections (Barrangou et al., 2007; Brouns et al., 2008; Marraffini and Sontheimer, 2008). Adaptation is achieved through the integration of small fragments called spacers acquired from invasive nucleic acids (Jackson et al., 2017; Lee and Sashital, 2022). These spacers provide the basis for RNA-guided interference leading to degradation of the foreign nucleic acid upon subsequent exposure (Marraffini, 2015). The CRISPR array consists of a leader sequence followed by direct repeats interspersing the acquired spacers, with addition of new spacers occurring almost invariably at the junction of the leader and the first repeat (Arslan et al., 2014; Bolotin et al., 2005; Mojica et al., 2005; Nuñez et al., 2015a, 2015b; Pourcel et al., 2005; Rollie et al., 2015; Yosef et al., 2012).

Target binding and degradation of the invading nucleic acids is preceded by recognition of the PAM (protospacer adjacent motif) located adjacent to the target sequence (Mojica et al., 2009; Redding et al., 2015; Sashital et al., 2012; Semenova et al., 2011; Sternberg et al., 2014; Xue et al., 2017). Because PAM recognition is required for interference, spacers acquired during adaptation must be selected from “prespacers” that contain a PAM sequence (Hu et al., 2021; Wang et al., 2015; Xiao et al., 2017). Importantly, the PAM is not incorporated into the CRISPR array, preventing “self”-targeting by the interference machinery and necessitating precise removal of the PAM before integration. Integration must take place in a polarized fashion with the non-PAM end and PAM end inserted at the leader-repeat junction on the plus strand and the repeat-spacer junction on the minus strand, respectively, to ensure generation of a crRNA complementary to the target (Jackson et al., 2017; Lee and Sashital, 2022).

Adaptation is mediated by the Cas1-Cas2 integration complex. Cas1 and Cas2 proteins form a heterohexameric complex with two distal Cas1 dimers flanking a Cas2 dimer (Fagerlund et al., 2017; Hu et al., 2021; Lee et al., 2019; Nuñez et al., 2014; Rollins et al., 2017; Xiao et al., 2017). While *cas1* and *cas2* are nearly universally conserved in diverse CRISPR-Cas systems, additional factors are required for adaptation in various sub-types (Makarova et al., 2020). Cas4 is the most common ancillary protein involved in adaptation, encoded by type I, II, and V CRISPR-Cas systems (Hudaiberdiev et al., 2017). While Cas4 usually exists as a standalone protein, some systems (type I-G, V-B) encode Cas4/1 fusion proteins, providing evidence of a role for Cas4 in adaptation. *In vivo* studies have demonstrated that Cas4 is essential for efficient spacer acquisition (Li et al., 2014), ensuring acquisition of spacers with correct PAM sequences (Almendros et al., 2019; Dixit et al., 2021; Kieper et al., 2018, 2021; Zhang et al., 2012) and in the correct orientation (Shiimori et al., 2018). Our previous *in vitro* studies of the type I-C system from *Alkalihalobacillus halodurans* (previously *Bacillus halodurans*) revealed that Cas4 acts as an endonuclease and cleaves specifically and precisely upstream of PAM sequences within 3′ overhangs of prespacers (Lee et al., 2018, 2019). Cas4 forms a higher order complex with Cas1-Cas2 in the presence of a dsDNA substrate, and is only activated as an endonuclease within this complex (Lee et al., 2019). Recent cryo-electron microscopy (cryo-EM) structures of an adaptation complex from the type I-G system demonstrated that Cas4/1 fusion proteins enable polarized integration through activation of Cas4 cleavage following the first integration event into the CRISPR array (Hu et al., 2021). However, it remains unclear how systems with standalone Cas4 and Cas1 proteins activate Cas4 as an endonuclease for PAM cleavage and enable polarized integration of new spacers.

Here we used cryo-EM to solve high resolution structures of the type I-C Cas4-Cas1-Cas2 complex in the presence of multiple substrates designed to mimic different stages of prespacer processing and integration. Our structural and biochemical results reveal how Cas4 recognizes the PAM with high specificity and sequesters the overhang away from the Cas1 active site. Flexible binding of Cas4 at the non-PAM end slows down trimming by non-specific exonucleases. Unlike in the type I-G Cas4/1 fusion-containing system, we do not observe enhanced endonuclease activation of standalone Cas4 following integration of the non-PAM end. Instead, tight binding of Cas4 to the PAM end of the prespacer and delayed PAM cleavage ensures insertion of the spacer in the correct orientation. Overall, this work demonstrates how Cas4-Cas1-Cas2 achieves processing and integration of functional spacers in the type I-C system.

## Results

### Cryo-EM structure of the Cas4-Cas1-Cas2 complex

We previously showed in the *A. halodurans* type I-C system that Cas4 is required for prespacer processing precisely upstream of the PAM (Lee et al., 2018, 2019) (**Figure 1A-C**). Efficient prespacer processing requires both Cas1 and Cas2, and is dependent on the Cas4 active site, but not on the Cas1 active site (**Figure 1C**). To investigate how Cas4 is activated within this complex, we solved a 3.4 Å structure of the type I-C Cas4-Cas1-Cas2-prespacer complex using cryo-EM (**Figures 1D, S1A, S1C, S1H and S2**). We initially reconstituted a complex bound to a prespacer substrate comprising a 22-base pair (bp) duplex with unprocessed 15 nucleotide (nt) 3′ overhangs, each containing the 5′-GAA-3′ PAM sequence beginning at the seventh position of the overhangs (**Figures 1E and S1A-C**). Consistent with our previous negative stain reconstruction and the recent cryo-EM structure of the type I-G Cas4/1-Cas2 complex (Hu et al., 2021; Lee et al., 2019), the cryo-EM structure reveals the typical butterfly shape of Cas1-Cas2. Two Cas1 dimers flank a Cas2 dimer in the middle and a single Cas4 subunit is associated with the wing tip on one end of each Cas1 dimer, resulting in a stoichiometry of Cas4_2_:Cas1_4_:Cas2_2_ (**Figures 1D and 1E**).

**Figure 1:**
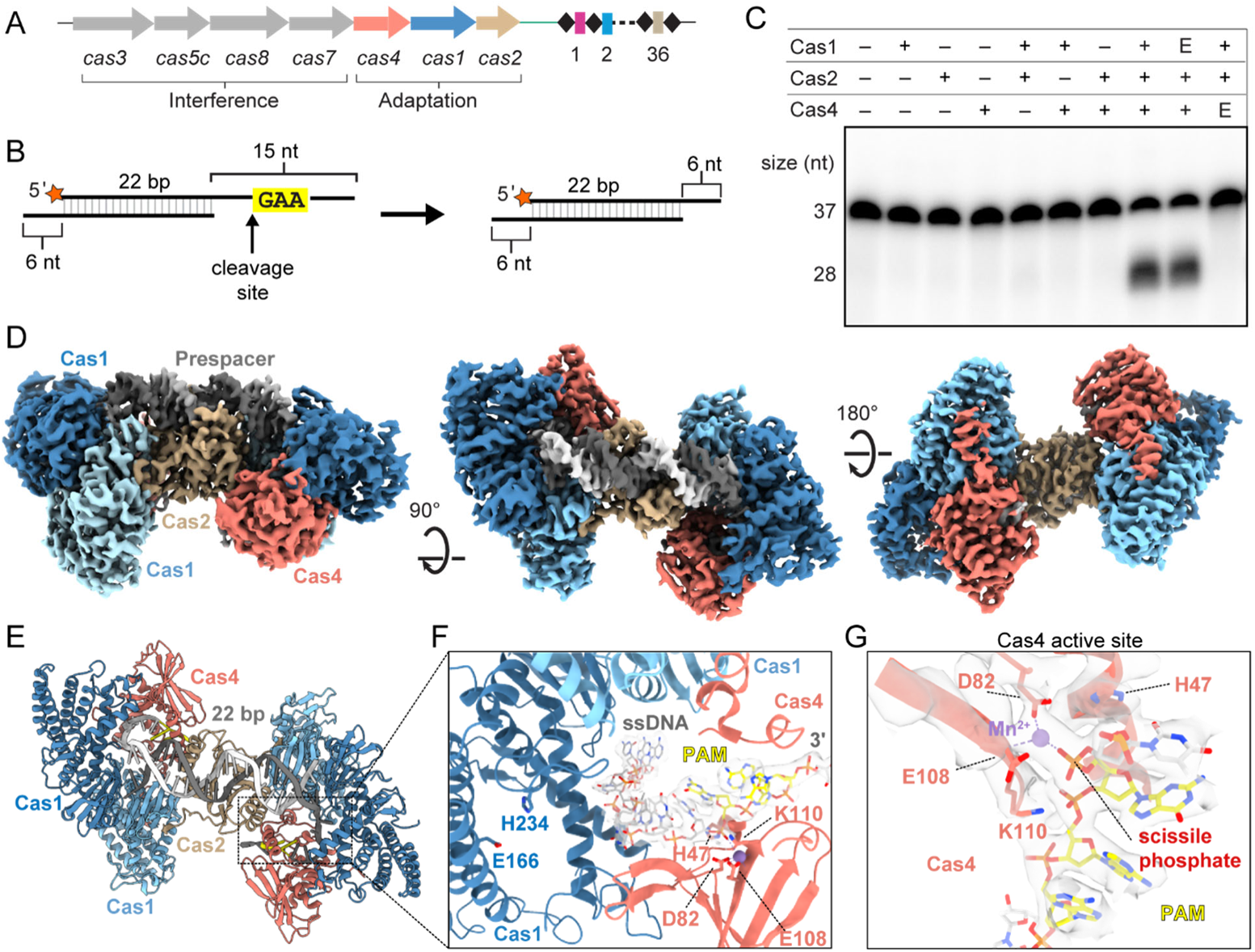
also see Figure S1, S2 and S3: Cryo-EM structure of the type I-C Cas4-Cas1-Cas2 complex. (A) Overview of *cas* genes and CRISPR locus of type I-C system in *Alkalihalobacillus halodurans. cas* genes are shown as arrows with the gene products involved in adaptation or interference indicated. Spacers and repeats are shown as colored rectangles and black diamonds respectively. (B) Schematic of prespacer cleavage assay. Cleavage site upstream of PAM is indicated with an arrow, radiolabel is indicated with an orange star. (C) Cleavage assay with substrate shown in (B). E indicates a Cas1 active site mutant E166A or Cas4 active site mutant E108A. Sizes for substrate (37 nt) and product (28 nt) are indicated. (D) Cryo-EM reconstruction of Cas4-Cas1-Cas2 complex with PAM/PAM substrate (Figure S1B). Subunits are labeled and colored as follows: dark blue, active Cas1 subunits; light sky blue, inactive Cas1 subunits; tan, Cas2 dimer; salmon, Cas4; dark gray and white, prespacer strands; yellow, PAM. (E) Structural model of the Cas4-Cas1-Cas2 PAM/PAM complex, colored as in (D). (F) Close up view of cryo-EM density (white) for single-stranded overhang. PAM residues are shown in yellow. Active site residues of Cas1 and Cas4 are labeled. (G) Close-up of Cas4 active site. The DEK motif residues (D82, E108 and K110) and His47 are shown as sticks. Cryo-EM density is shown for the coordinated Mn^2+^ ion, portions of Cas4, and the DNA in proximity to the active site.

Cryo-EM density for the Cas4 subunits was sufficiently high resolution to build a structural model for a type I-C Cas4 protein using an AlphaFold predicted structural model for guidance (**Figure S3A**) (Jumper et al., 2021; Mirdita et al., 2022; Varadi et al., 2022). Type I-C Cas4 adopts an overall fold similar to previously determined structures of type I-A, I-B and I-G Cas4 proteins (Hu et al., 2021; Lemak et al., 2013, 2014). The C-terminus of type I-C Cas4 extends further than in the other structures, adopting an α-helix (**Figures 1E and S3B-C**). While the Cas4 subunits mostly interact with the active subunits of Cas1, the Cas4 C-terminal helix mediates an additional interaction with the inactive Cas1 subunit (**Figure S3C**). The N-terminus of type I-C Cas4 is located on the opposite side of the protein in comparison to type I-A and I-B, but in a similar location to type I-G (**Figure S3B**). The N-terminus of Cas4 fits into a positively charged groove near the Cas1 active site, potentially blocking access to a prespacer overhang in the presence of Cas4 (**Figure S3D**). Consistently, our previous biochemical results indicated that Cas4 prevents premature integration of unprocessed prespacers by Cas1 (Lee et al., 2018).

The cryo-EM map contains continuous density for the single-stranded overhang through the Cas4 active site (**Figure 1F**). Conserved active site residues D82 and E108 are in proximity of a coordinated Mn^2+^ ion that is positioned to enable cleavage upstream of the 5′-G_29_-A_30_-A_31_-3′ PAM sequence (**Figures 1F and 1G**). While the three residues of the DEK RecB motif (D82, E108 and K110) are positioned similarly to a recent AdnAB structure (Jia et al., 2019), a conserved histidine (H47) that was observed to coordinate divalent ions in both AdnAB and the type I-G Cas4/1 structures (Hu et al., 2021; Jia et al., 2019), is unambiguously pointed away from the Mn^2+^ ion in the type I-C structure (**Figure 1G**). In addition, density for the scissile phosphate is consistent with an intact phosphodiester bond. Together, these observations suggest that the Cas4 active site structure was captured in an inactive conformation and that cleavage did not occur during complex preparation, potentially due to the formation of the complex on ice, at a suboptimal temperature for cleavage.

### Activation of Cas4 endonuclease activity

While type I-A and I-B Cas4 proteins have been demonstrated to have nonspecific nuclease activities in the absence of Cas1 and Cas2 (Dixit et al., 2021; Lemak et al., 2013, 2014), type I-C Cas4 functions as a sequence specific endonuclease that is only activated in the presence of Cas1 and Cas2 (Lee et al., 2018) (**Figure 1C**). To understand this activation, we examined features of Cas4 that interact with either Cas1 or Cas2. A striking feature of the type I-C Cas4 structure is the C-terminal helix, which is unique to type I-C, based on comparison of known Cas4 structures and multiple sequence alignment of Cas4 protein sequences from all Cas4-containing CRISPR-Cas subtypes (**Figure 2B, S3B**). Besides interacting with Cas1, Cas4 also interacts with Cas2 via its α4 that forms close contacts with α2 of Cas2 (**Figure 2C**), similar to an interaction observed in the type I-G Cas4/1-Cas2 structure. Notably, the trajectory of the substrate DNA passes through a narrow hole created, in part, by Cas4 α4, suggesting that the position of this helix is important for accommodation of the DNA. Together, these interactions may explain why Cas4 cleavage activation requires both Cas1 and Cas2 (**Figure 1C**).

**Figure 2:**
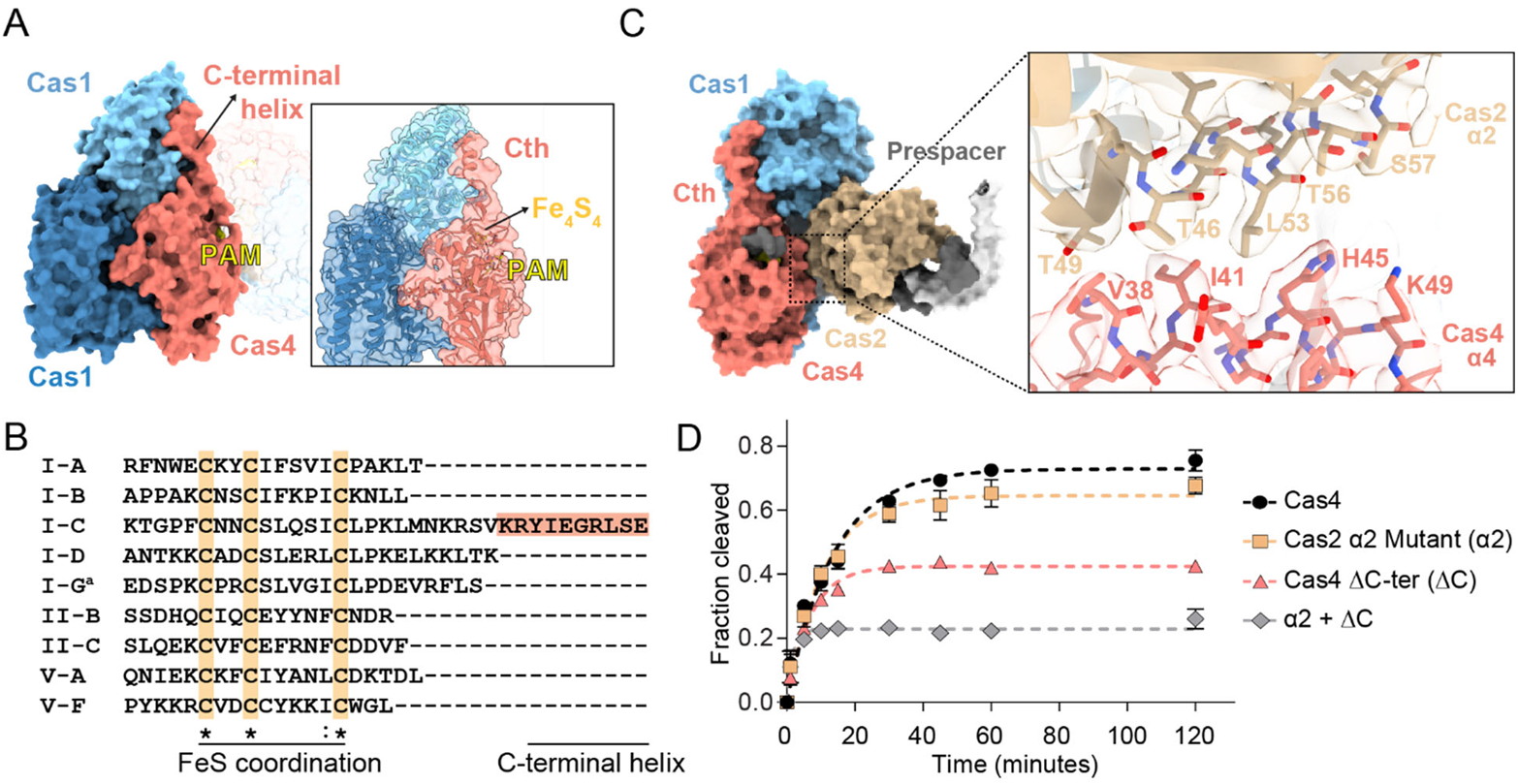
also see Figure S4: Activation of Cas4 endonuclease activity. (A) Side surface view of Cas4-Cas1-Cas2 PAM/PAM complex with C-terminal helix and PAM labeled. Inset shows structural model with transparent surface. Iron-sulfur cluster and PAM are labeled. (B) Clustal Omega multiple sequence alignment for the C-terminus of representative Cas4 sequences from Cas4-containing CRISPR-Cas systems. ′a′ indicates the type I-G sequence, which is a Cas4/1 fusion protein. The sequence was truncated at the end of the Cas4 domain based on structural analysis (Hu et al., 2021). Cas4 sequences were used from the following organisms: *Saccharolobus solfataricus* (I-A); *Pyrobaculum calidifontis* (I-B); *Alkalihalobacillus halodurans* (I-C); *Synechocystis sp.* (I-D); *Geobacter sulfurreducens* (I-G); *Francisella tularensis subsp. novicida* (II-B); *Micrarachaeum acidiphilum* (II-C); *Francisella tularensis subsp. novicida* (V-A); *Uncultured archaeon* (V-F). Type V-B was omitted from the analysis due to the presence of a Cas4/1 fusion, for which the end of the Cas4 domain cannot be precisely determined. (C) Surface view of one lobe of the complex highlighting interaction between α2 of Cas2 and α4 of Cas1. The close-up shows the structural model fit in the cryo-EM density for the two helices. (D) Quantification of fraction cleaved for cleavage assay with individual Cas4 (ΔCth) or Cas2 α2 mutant (alanine substitutions of Thr46, Thr49, Leu53, Thr56 and Ser57) or a combination of the two. The assay was performed at final concentrations of 250 nM Cas1, 250 nM Cas2 and 2 µM Cas4. The data points were fit to a one phase exponential association. Polyacrylamide gel for the cleavage assay is shown in Figure S4C. The average of three replicates is shown, error bars represent standard deviation.

To determine the importance of these interactions for Cas4 cleavage, we designed a Cas4 C-terminal helix deletion (ΔCth) and a Cas2 α2 mutant with Thr46, Thr49, Leu53, Thr56 and Ser57 residues participating in the Cas4-Cas2 interface substituted with alanine (**Figures 2D, S4A-C**). Cleavage activity was severely diminished upon deletion of the C-terminal helix. In contrast, Cas2 α2 substitutions did not have a substantial effect on cleavage, potentially due to the relatively conservative mutations of threonine or serine to alanine. Interestingly, a combination of the Cas2 α2 substitutions and Cas4 ΔCth lead to severe loss in cleavage activity of Cas4. It is possible that this loss of cleavage activity is due to a decrease in Cas4 affinity for the Cas1-Cas2 complex, a loss of important interactions that serve to activate Cas4 nuclease activity, or a combination of the two. We performed cleavage using the two mutants either alone or in combination at increasing Cas4 concentration with limited Cas1-Cas2 concentrations (**Figures S4A and S4B**). We did not observe statistically significant differences in cleavage activity at increasing Cas4 variant concentrations, suggesting that Cas4 cleavage is impaired even when Cas4 concentration is saturating. Overall, the additive cleavage defects of these Cas4 variants suggests that interactions with both Cas1 and Cas2 are important for activating Cas4 as an endonuclease.

### PAM recognition by Cas4

Cas4 recognizes a GAA PAM sequence positioned within a tight groove that guides the single-stranded overhang through the RecB nuclease domain (**Figure 3A**). Within the PAM, the two adenine bases adopt a syn conformation, with rotation of the N-glycosidic bond resulting in placement of the adenine bases above the deoxyribose. Polar and non-polar residues including Asn37, Thr40, Glu119, Gln123, Leu195 and Ile28 aid in shape recognition of the PAM nucleotides and formation of the tight PAM recognition channel (**Figure 3B**). Each nucleotide of the PAM potentially participates in hydrogen bonding interactions (**Figures 3B and 3C**). The first nucleotide of the PAM, G29, is in proximity of polar residues His17 and Gln44, with potential formation of a hydrogen bond between His17 and N7 of the G29 purine ring. The second nucleotide of the PAM, A30, is recognized by Ser194 that forms two potential hydrogen bonds with N6 and N7 of the purine ring respectively (**Figures 3B and 3C**). The PAM is further specified by the last nucleotide A31 that forms two bidentate interactions, which is facilitated by the syn conformation of the adenine base. N1 and N6 of the adenine are in proximity to form hydrogen bonds with Gln16 and N7 and N6 with Gln24 (**Figures 3B and 3C**). Two hydrophobic residues, Phe20 and Trp34, sit across the phosphodiester backbone. Phe20 is placed after A31 and Trp34 lies between G29 and A30 (**Figures 3B and 3C**), providing stabilizing stacking interactions for the two purine bases that would be minimized in case of pyrimidines.

**Figure 3:**
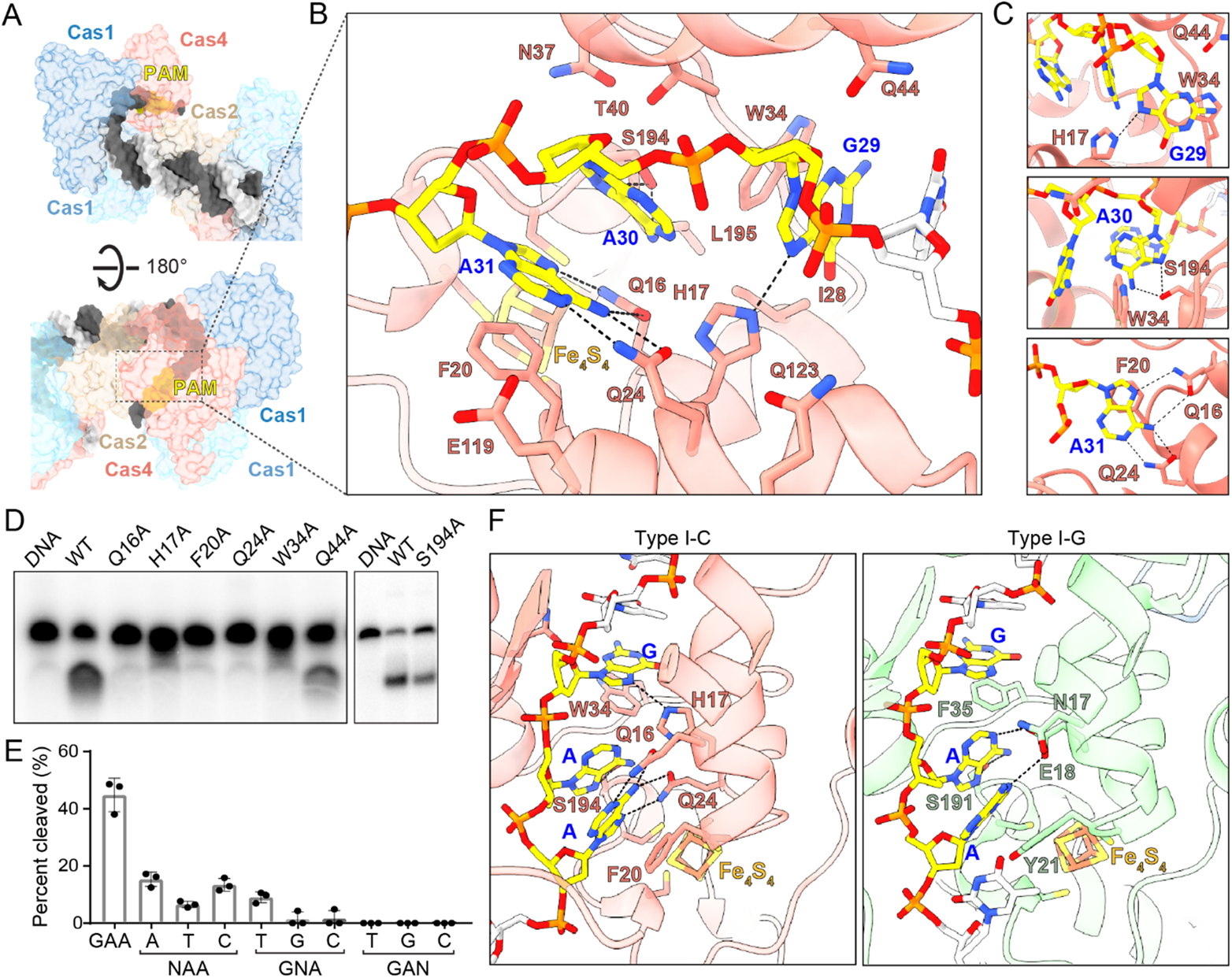
PAM recognition by Cas4. (A) Surface view of single-stranded overhang travelling to Cas4 PAM recognition pocket. (B) Specific and non-specific interactions between Cas4 residues (labeled in salmon with black outline) and PAM nucleotides (labeled in blue). Potential hydrogen bonds are shown as dashed lines. (C) Close-up view of potential hydrogen bonds formed by individual PAM nucleotides. (D) Polyacrylamide gel showing cleavage assay with PAM recognition mutants. The assay was performed with final concentrations of 250 nM Cas1, 250 nM Cas2 and 500 nM Cas4 and quenched at 30 minutes. (E) Quantification of cleavage assay for saturation mutagenesis of each nucleotide in the PAM sequence, performed as in (D). N indicates the position that was mutated. The average of three replicates is shown, error bars represent standard deviation and the three data points are plotted as dots. (F) Comparison of I-C and I-G Cas4 showing specific and stacking interactions for PAM recognition.

To test the importance of these interactions on PAM recognition and prespacer processing, we introduced alanine substitution at each of these potential interacting residues (**Figure 3D**). All substitutions ablated cleavage activity, with the exception of Q44A and S194A, which only partially reduced cleavage. Gln44 is located within helix α4, on the opposite face of the helix that interacts with helix α2 of Cas2 (**Figure 2C**). The minimal defect in cleavage for the Q44A substitution suggests that Gln44 is not involved in PAM recognition but potentially facilitates widening of the PAM recognition channel to allow correct positioning of the PAM. Partial reduction in cleavage with S194A further indicates that shape recognition mediated by neighboring residues, rather than hydrogen bonding, plays the major role in recognition of the second nucleotide of PAM, A30 (**Figure 3D**). Additionally, we performed saturation mutagenesis to validate the importance of single nucleotides in the PAM sequence (**Figure 3E**). We observed that cleavage is severely compromised when we replaced the first or second nucleotides of the PAM, G29 or A30, and is completely lost when the last nucleotide of the PAM A31 is changed. Together, these results strongly suggest that the bidentate recognition of A31 by Gln16 and Gln24 is essential for specific PAM recognition by Cas4 (**Figure 3C-E**).

We compared the PAM recognition motifs for type I-C Cas4-Cas1-Cas2 and type I-G Cas4/1-Cas2 complexes (Hu et al., 2021) to understand differences that might affect the overall specificity of the two systems (**Figure 3F**). The type I-G adaptation complex lacks specificity at the first position of the PAM, recognizing a 5′-NAA-3′ PAM sequence, in contrast to the relative specificity for 5′-GAA-3′ observed in type I-C (Almendros et al., 2019; Rao et al., 2017). Consistently, the equivalent residue to His17 in I-C is Glu18 in I-G, which is positioned slightly further from the guanine, but could form a hydrogen bond with N1 of either adenine in the PAM upon protonation of the Glu18 carboxylate (**Figure 3F**). Similarly, Trp34, which stacks with the guanine in I-C, is replaced with Phe35 in I-G. This substitution may decrease the stacking interaction energy and possibly reduce specificity for a purine at this position of the PAM in I-G (Rutledge et al., 2006).

Recognition of the third position of the PAM also differs between the two structures. While the adenine in the type I-C structure is stabilized in the syn conformation based on a dual-bidentate interaction with Gln16 and Gln24, the type I-G structure positions the equivalent adenine in an anti conformation, enabling the aforementioned potential hydrogen bonding interaction with Glu18 (**Figure 3F**). Gln16 of I-C is substituted with Asn17 in I-G. This shorter side chain and slightly further positioning prevents recognition of the adenine via specific hydrogen bonding interactions that are observed in type I-C. Gln24 of I-C is substituted with Leu25 in I-G, which mediates van der Waals contacts with the adenine for shape readout but does not enable specific hydrogen bonding. Overall, these differences suggest that Cas4 may confer higher specificity for a GAA PAM in the type I-C system.

### Cas4-Cas1-Cas2 complex specifies the length of the prespacer duplex and overhang

In addition to the 22 bp duplex prespacer, we also solved the cryo-EM structure of a complex with a 24 bp duplex with 3′ overhangs containing PAMs starting at the sixth position (**Figures S1A, S1B, S1G, S1H and S5A-G**). The reconstruction has weaker density for one of the active Cas1 subunits, while density for the partner subunit is stronger, suggesting conformational heterogeneity (**Figures S5H and S5I**). Additionally, density for the prespacer corresponds to a 22 bp duplex, suggesting unwinding of the last base pair at either end of the duplex, and creating a 6 nt overhang up to the PAM sequence (**Figure S5I**). A conserved tyrosine in Cas1 plays a role in defining the duplex in potential prespacers in other systems like type I-E (Nuñez et al., 2015a). Tyr49 is positioned similarly in the Cas4-Cas1-Cas2 structure, suggesting a similar role for duplex definition in type I-C systems (**Figure 4A**).

**Figure 4:**
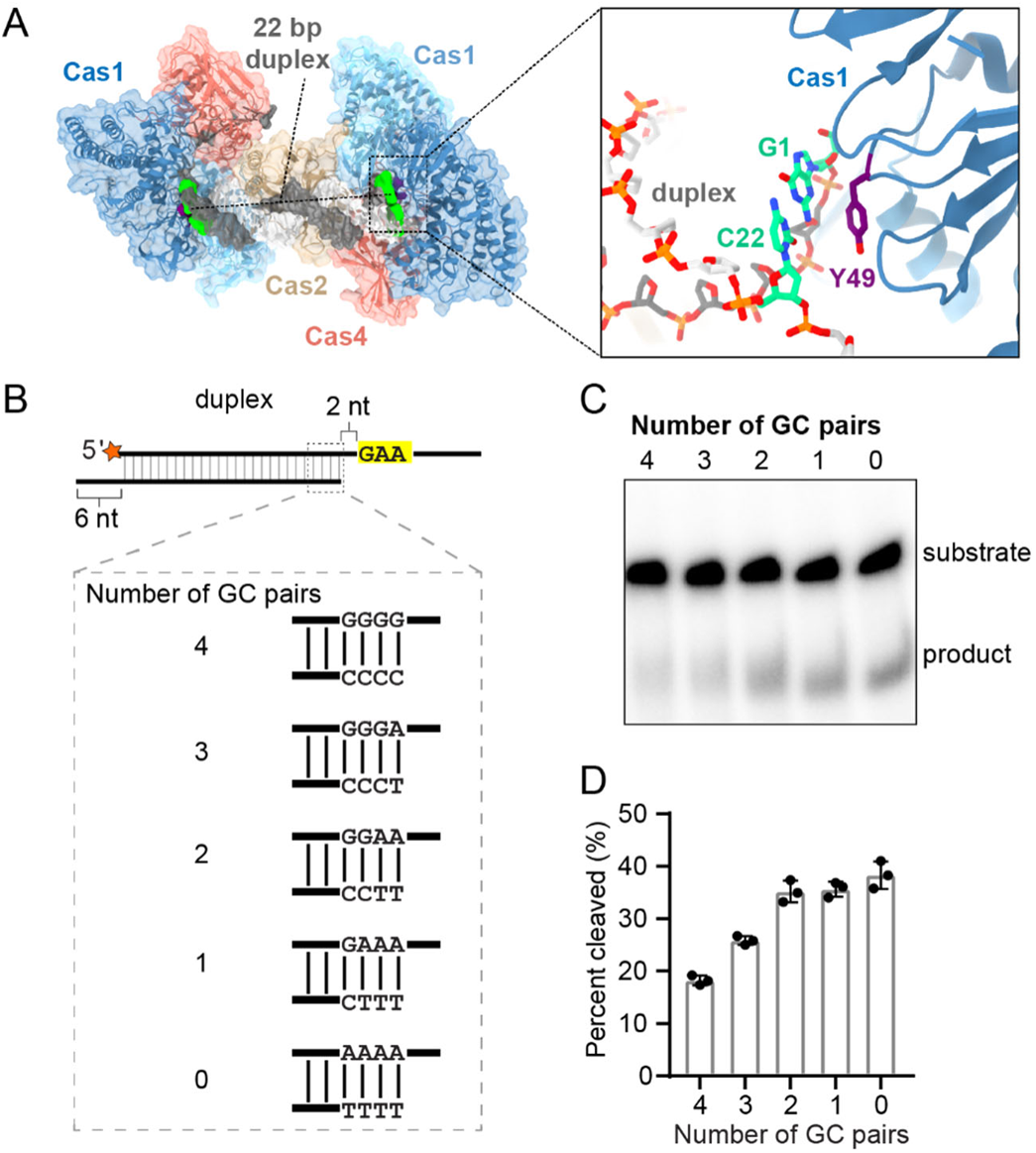
also see Figure S1 and S5: Cas4-Cas1-Cas2 complex specifies the length of the prespacer duplex and overhang. (A) Surface view of Cas4-Cas1-Cas2 PAM/PAM structural model highlighting 22 bp duplex and Tyr49 from both active Cas1 subunits. Close up view of Tyr49 (purple) interaction with the last base pair (green) of the 22 bp duplex of the prespacer is shown. (B) Panel of substrates used in unwinding and processing assays. Substrates contain between zero and four GC pairs at the end of the 26 bp duplex. The 5′-GAA-3′ PAM starts 2 nt after the end of the 26 bp duplex. Radiolabel is indicated with a star. (C) Representative denaturing polyacrylamide gel of cleavage of substrates shown in (B). The assay was performed with final concentrations of 250 nM Cas1, 250 nM Cas2 and 500 nM Cas4 and quenched at 30 minutes. (D) Quantification of cleavage assay shown in (C). The average of three replicates is shown, error bars represent standard deviation and the three data points are plotted as dots.

The two Cas4-Cas1-Cas2 structures indicate that a 6 nt single-stranded overhang is necessary to stretch the PAM from the end of the duplex to the Cas4 PAM recognition motif (**Figure 1F, S5I**). We previously observed that prespacer substrates with 4 nt between the duplex and the PAM were cleaved as efficiently as prespacers with 6 nt between the duplex and the PAM (Lee et al., 2019). Combined with our structural observations, these results imply that Cas4-Cas1-Cas2 can unwind multiple base pairs to access the single-stranded PAM. To test this further, we analyzed Cas4-dependent cleavage of a prespacer substrate containing a 26 bp duplex with 2 nt between the end of the duplex and the PAM. To determine the effect of unwinding, we varied the G-C content of the terminal 4 base pairs (**Figure 4B**). We observed a linear correlation between the number of terminal A-T base pairs and the amount of cleavage, suggesting that substrates that can be more readily unwound due to a higher number of A-T pairs result in better PAM accessibility (**Figures 4C and 4D**). Together, these results indicate that the Cas4-Cas1-Cas2 complex can unwind the ends of duplexes to allow Cas4 to gain access to the single stranded PAM region.

### Cas4 slows down exonucleolytic trimming of the non-PAM end

Cas4 cleavage is highly sequence specific and only processes the PAM end of the prespacer (Lee et al., 2019). The non-PAM end of the prespacer is likely trimmed by cellular non-specific exonucleases, similar to other type I systems where DnaQ-like exonucleases have been implicated in prespacer processing (Hu et al., 2021; Kieper et al., 2021; Kim et al., 2020; Ramachandran et al., 2020). To simulate the Cas4-Cas1-Cas2 complex before and after trimming of the non-PAM end, we solved cryo-EM structures of Cas4-Cas1-Cas2 bound to either a PAM/NoPAM (**Figures 5A, S1A, S1B, S1D, S1H and S6**) substrate or a PAM/processed substrate (**Figures 5B, S1A, S1B, S1E, S1H and S7**). Both substrates contained a 22-bp duplex with one 15-nt overhang containing a PAM and either another 15-nt overhang without a PAM (PAM/NoPAM substrate) (**Figure 5A and S1B**), or a 6-nt overhang simulating a processed non-PAM end (PAM/processed substrate) (**Figure 5B and S1B**).

**Figure 5:**
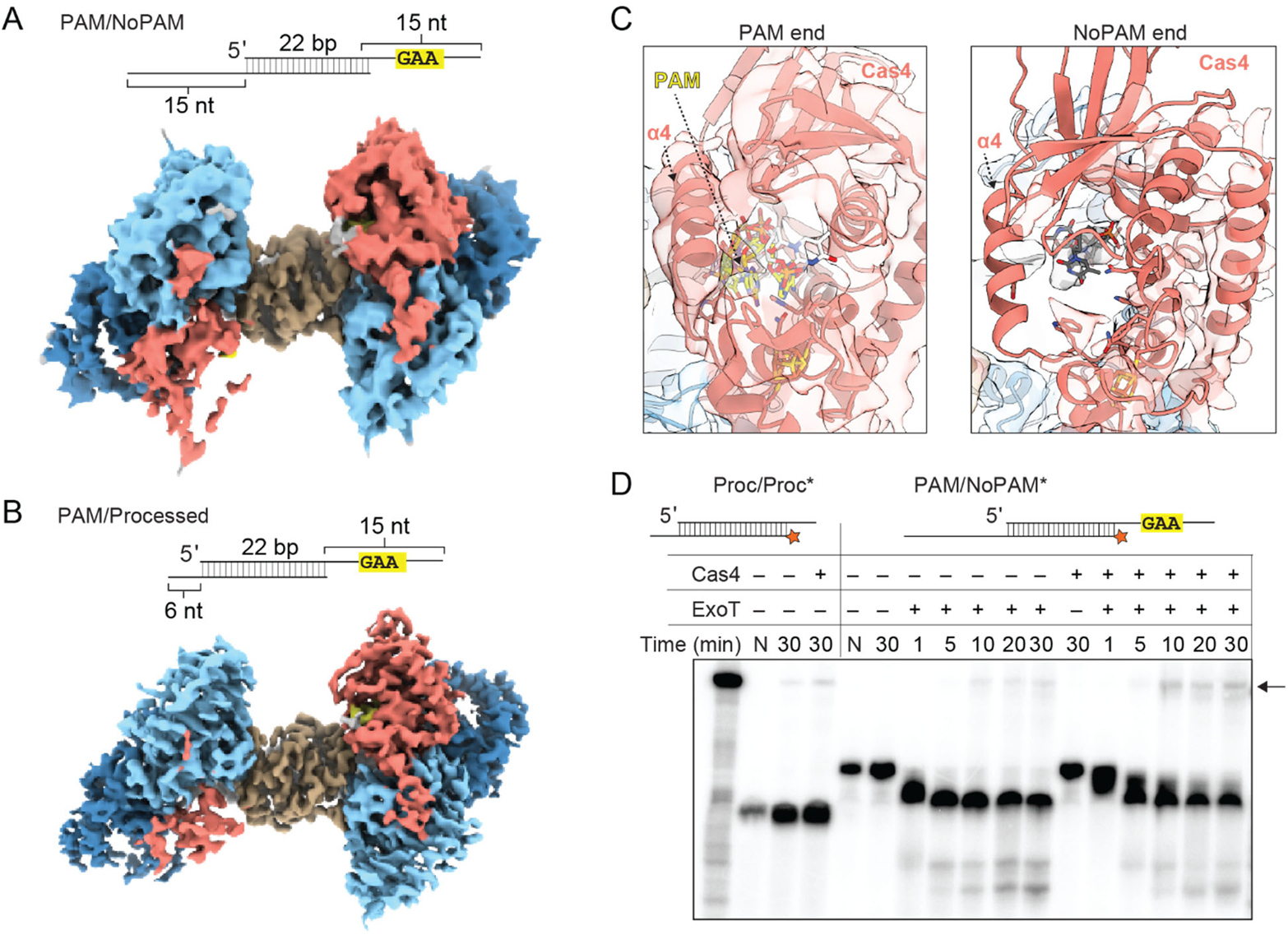
also see Figure S1, S6, S7 and S8: Cas4 slows down exonucleolytic trimming of the non-PAM end. (A) Substrate schematic and cryo-EM reconstruction of Cas4-Cas1-Cas2 PAM/NoPAM complex. (B) Substrate schematic and cryo-EM reconstruction of Cas4-Cas1-Cas2 PAM/processed complex. (C) Densities for Cas4 at PAM and No-PAM ends of PAM/NoPAM complex shown in (A). PAM is colored in yellow and labeled. (D) Denaturing polyacrylamide gel of the prespacer trimming and integration assays using ExoT exonuclease. The assay was performed with final concentrations of 100 nM Cas1, 50 nM Cas2 and 50 nM Cas4. Lanes with only substrate are labeled N. All other lanes have Cas1 and Cas2. Presence or absence of Cas4 and ExoT is indicated. Black arrow indicates integration products.

Single-particle analysis of these complexes yielded 3.9 Å and 3.3 Å reconstructions for the PAM/NoPAM and PAM/processed complexes, respectively (**Figures 5A and 5B**). For both maps, we observed only partial density for one of the two Cas4 subunits, resulting in asymmetrical reconstructions that we have previously observed by negative stain (Lee et al., 2019). Because Cas4 specifically binds to a PAM containing sequence, we concluded that the lobe with stronger density for Cas4 is the PAM end and the lobe with partial density is the non-PAM end of the substrate. We did not observe any significant differences in Cas4 conformation on the PAM end between the PAM/PAM and the PAM/NoPAM or PAM/processed complexes.

On the non-PAM end, we observed stronger density for a second Cas4 subunit when the longer 3′-overhang was present in the PAM/NoPAM substrate (**Figure 5A**). In contrast, the PAM/processed complex nearly completely lacked Cas4 density (**Figure 5B**). In the PAM/NoPAM reconstruction, density is present for the iron-sulfur cluster domain and C-terminal helix (**Figure 5A**). However, the density is weaker in regions that do not directly contact one of the Cas1 subunits. Notably, we observed a complete lack of density for α4 of Cas4 that typically interacts with a Cas2 α2 on the non-PAM end (**Figure 5C**). This lack of density suggests that α4 is conformationally flexible and does not interact stably with Cas2 α2 in the absence of PAM binding.

Similar to the PAM end, density for single-stranded DNA on the non-PAM end is traceable from the end of the duplex toward the Cas4 active site when the 15 nt overhang was present in the PAM/NoPAM substrate. However, no density is present past residue 28 of the DNA, consistent with a lack of stabilizing interactions with Cas4 in the absence of a PAM (**Figure 8A**). In the presence of the 6 nt overhang in the PAM/processed substrate, we could trace only 2 nt of ssDNA overhang density (**Figure S8B**). Overall, these structures suggest that longer DNA overhangs are necessary to anchor Cas4 to the Cas1-Cas2 complex, although Cas4 associates most stably when a PAM is present in the overhang.

Based on these structural observations, we hypothesized that Cas4 remains partially associated with Cas1-Cas2 in the presence of long 3′-overhangs and may affect trimming of the non-PAM end by cellular exonucleases. To test this, we used a commercially available DnaQ-like exonuclease, ExoT, to trim the non-PAM end of the PAM/NoPAM substrate in the presence of Cas1-Cas2 or Cas4-Cas1-Cas2 and a DNA substrate containing a leader-repeat-spacer to mimic a CRISPR array for integration (**Figures 5D and S8C**). ExoT trimming of the non-PAM end of the Cas1-Cas2-bound prespacer produced products of similar length to a pre-processed prespacer control, although trimming was slowed substantially in the presence of Cas4 (**Figure 5D**). The trimmed products were integrated into the CRISPR DNA, producing an integration product of similar size to the pre-processed control, with a slight enhancement of integration observed in the presence of Cas4. Overall, these results suggest that unstable association of Cas4 at the non-PAM end offers partial protection from exonucleolytic trimming, slowing down processing of the non-PAM end.

### Cas4-Cas1-Cas2 mediates polarized integration

Functional spacer acquisition requires integration of the non-PAM end of the prespacer at the leader-repeat junction on the positive strand (hereafter leader site) and the PAM end at the repeat-spacer junction on the negative strand (hereafter spacer site) (**Figure S8C**). Importantly, Cas4 association with Cas1-Cas2 is mutually exclusive with prespacer integration into the CRISPR array, as observed previously in our negative stain reconstructions and in type I-G cryo-EM structures (Hu et al., 2021; Lee et al., 2019). Cas4 dissociates from the non-PAM end, allowing integration at the leader site. A recent 5.8 Å structure of type I-G Cas4/1-Cas2 bound to a half-site intermediate (HSI) showed that the Cas4 domain remains associated with the PAM end of the prespacer following this integration event (Hu et al., 2021). The structure suggested that the adjacent Cas1 domain interacts with the repeat, although the low resolution of the reconstruction limited interpretation of this interaction.

To determine how the repeat interacts within the type I-C adaptation complex, we solved a 4 Å cryo-EM structure of the Cas4-Cas1-Cas2 complex bound to a half-site intermediate (**Figures 6A, 6B, S1A, S1B, S1F, S1H and S9**). We used the Cas1 active site mutant E166A to prevent disintegration of the prespacer strand integrated at the leader site of the HSI mimic. Strong density for the CRISPR DNA could be observed in the reconstruction (**Figures 6A and 6B**). The DNA helix bends sharply near the leader-repeat junction with another slight bend near the Cas4-Cas2 interface. The repeat spans the side of the complex opposite to the prespacer duplex. Unlike in the type I-G structure, where the repeat appears to contact the inactive Cas1 domain of the Cas4/1 subunit, in the type I-C structure, the repeat contacts the C-terminal helix of Cas4 (**Figure 6C**). Specifically, two arginine residues Arg207 and Arg211 in the Cas4 C-terminal helix point directly at the terminal sequence of the repeat, interdigitating within the minor groove of the repeat end of the CRISPR (**Figure 6C**). This may suggest a role of Cas4 in correctly orienting the complex onto the CRISPR array, positioning the PAM end of the prespacer close to the repeat-spacer site for integration following PAM cleavage and Cas4 dissociation.

**Figure 6:**
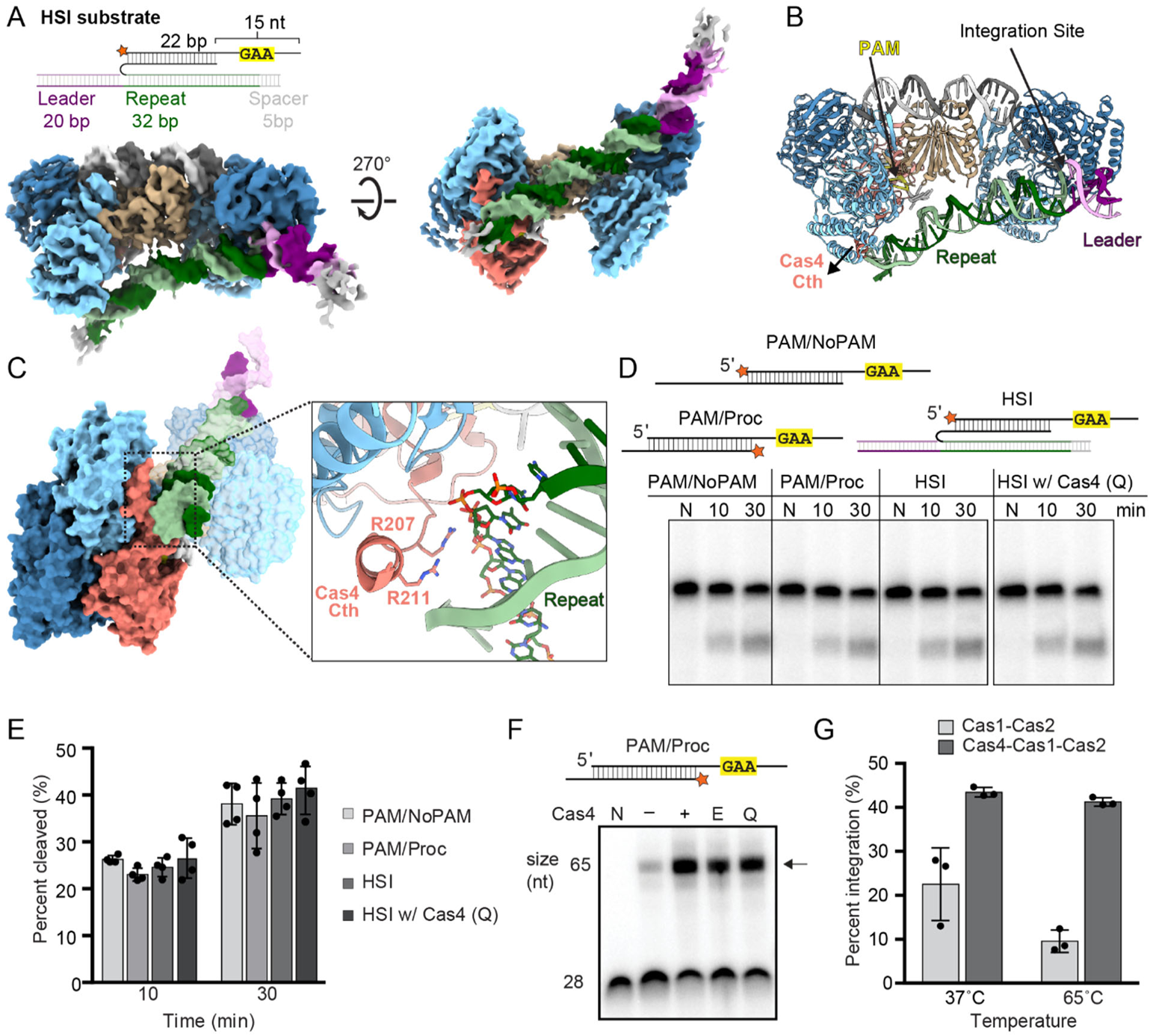
also see Figure S1, S9, S10 and S11: Cas4-Cas1-Cas2 mediates polarized integration. (A) Substrate schematic and cryo-EM reconstruction of Cas4-Cas1-Cas2 HSI complex. Protein subunits and prespacer strands are colored as in Figure 1D. Leader and repeat DNA are colored purple and green, respectively. (B) Structural model for HSI complex with integration site, PAM, leader, repeat and Cas4 C-terminal helix (Cth) labeled. (C) Side view of HSI complex with close-up view of Cas4 C-terminal interaction with the repeat DNA. (D) Representative denaturing polyacrylamide gel of cleavage assay with PAM/NoPAM, PAM/processed (PAM/Proc) and HSI substrates. Wild type Cas4 was used for PAM/NoPAM, PAM/processed and HSI samples, while the Cas4 repeat-interacting residue mutant (Q; Alanine substitutions of Lys206, Arg207, Lys210 and Arg211) was used for gel on right. The assay was performed with final concentrations of 100 nM Cas1, 100 nM Cas2 and 50 nM Cas4. Radiolabel is indicated with a star. Lanes for samples with only DNA are labeled N, or time points are labeled in minutes. (E) Quantification for percent cleaved in (D). The average of four replicates is shown, error bars represent standard deviation and the four data points are plotted as dots. (F) Representative denaturing polyacrylamide gels showing integration of the processed end in the absence and presence of WT Cas4, Cas4 active-site mutant (E108A) and Cas4 repeat-interacting residue mutant (Q). The assay was performed with final concentrations of 100 nM Cas1, 50 nM Cas2 and 50 nM Cas4 and quenched at 30 minutes. Lanes with only substrate are labeled N. All other lanes contained Cas1 and Cas2. Presence or absence of Cas4 is indicated. Black arrow indicates integration products. (G) Quantification of substrate integration in the presence and absence of Cas4 at different temperatures. The average of three replicates is shown, error bars represent standard deviation and the three data points are plotted as dots.

In the type I-G system, PAM cleavage by the Cas4 domain of Cas4/1 is activated following integration of the prespacer at the leader site, potentially due to the interaction between the repeat and the inactive Cas1 domain (Hu et al., 2021). Cleavage is followed by integration of the PAM end at the spacer site. Overall, this mechanism ensures insertion of a correctly oriented spacer. However, we have previously observed that type I-C Cas4 cleavage activity is not altered in the presence of a CRISPR array (Lee et al., 2019), suggesting that Cas4 is not activated through interactions with the CRISPR following integration of the non-PAM end. To further investigate whether type I-C Cas4 is activated following half-site integration, we compared Cas4 cleavage of the PAM end within an HSI substrate with cleavage of the PAM/NoPAM and PAM/processed substrates (**Figures 6D and 6E**). Similar to our previous results, we did not observe any significant differences in the fraction of substrate cleaved at various time points between the substrates, suggesting that Cas4 processing is not dependent on the integration of the non-PAM strand at the leader site. To test whether interactions between the Cas4 C-terminal helix and the repeat affect PAM-end processing following leader-site integration, we tested a quadruple mutant of Cas4 (Q) with the two C-terminal arginine residues and two nearby lysine residues (Lys206 and Lys210) substituted with alanine (**Figures 6D and 6E**). Mutations of these residues had no effect on PAM cleavage within the HSI substrate. Overall, these results strongly suggest that Cas4 is not activated through interactions with the repeat following non-PAM integration in the type I-C adaptation complex, in contrast to the type I-G system.

### Cas4 enhances integration of the prespacer non-PAM end

As noted above, in the exonuclease trimming assays, we observed slightly more integration in the presence of Cas4, despite observing slower exonucleolytic cleavage in this condition (**Figure 5D**). This led us to hypothesize that Cas4 may enhance integration at the leader site. To measure this effect more precisely, we used a PAM/processed substrate with the processed strand radiolabeled to measure integration of this strand at the leader site in the absence and presence of Cas4 (**Figures 6F and S8C**). We observed substantial enhancement of integration at multiple temperatures in the presence of Cas4 (**Figures 6F and G**), while Cas4 had no detectable effect on Cas1 disintegration activity (**Figure S10**). These observations suggest an additional role for Cas4 within the Cas4-Cas1-Cas2 complex in enhancing integration at the leader site.

We next tested whether enhanced integration is an effect of PAM cleavage in the presence of Cas4 or interactions between the Cas4 C-terminal helix and the repeat. However, neither a Cas4 active site mutant nor the quadruple Cas4 C-terminal mutant (Q) substantially affected the enhancement of integration (**Figure 6F**). Similarly, substrates containing phosphorothioate substitutions at the scissile phosphate that prevent PAM cleavage still displayed enhanced integration in the presence of Cas4 (**Figure S11**). Notably, a substrate containing processed 6-nt overhangs on both ends was not integrated well either in the presence or absence of Cas4 (**Figure S11B**). Overall, these results strongly suggest that Cas4 enhances integration of the processed end at the leader site due to its tight binding to a PAM-containing substrate rather than its PAM cleavage activity or its interaction with the CRISPR repeat.

## Discussion

Our structural and biochemical results provide a kinetic model for prespacer processing and integration by the type I-C Cas4-Cas1-Cas2 complex (**Figure 7**). During prespacer capture, Cas4 is essential for binding prespacers containing a PAM sequence (**Figure 7A**). These prespacers may be single-stranded or double-stranded fragments of degradation products, potentially of the AddAB complex (Ivančić-Bace et al., 2015; Kim et al., 2020; Levy et al., 2015; Modell et al., 2017). While Cas4 stably binds to the PAM end, it remains flexible on the non-PAM end, slowing trimming of this end to the proper length by cellular exonucleases (**Figures 7A and 7B**). Nevertheless, trimming of the non-PAM end gives rise to an asymmetric substrate based on the slow kinetics of Cas4 PAM cleavage. Following trimming, Cas4 dissociates from the non-PAM end (**Figures 7B and 7C**). Tight binding of Cas4 at the PAM end increases the likelihood of integration of the non-PAM end at the leader site, enabling rapid formation of a half site integration intermediate after exonucleolytic trimming (**Figures 7C and 7D**). Following PAM cleavage, Cas4 dissociates from the complex, allowing the mature PAM end to be integrated at the spacer site resulting in insertion of a polarized and functional spacer (**Figures 7E and 7F**).

**Figure 7:**
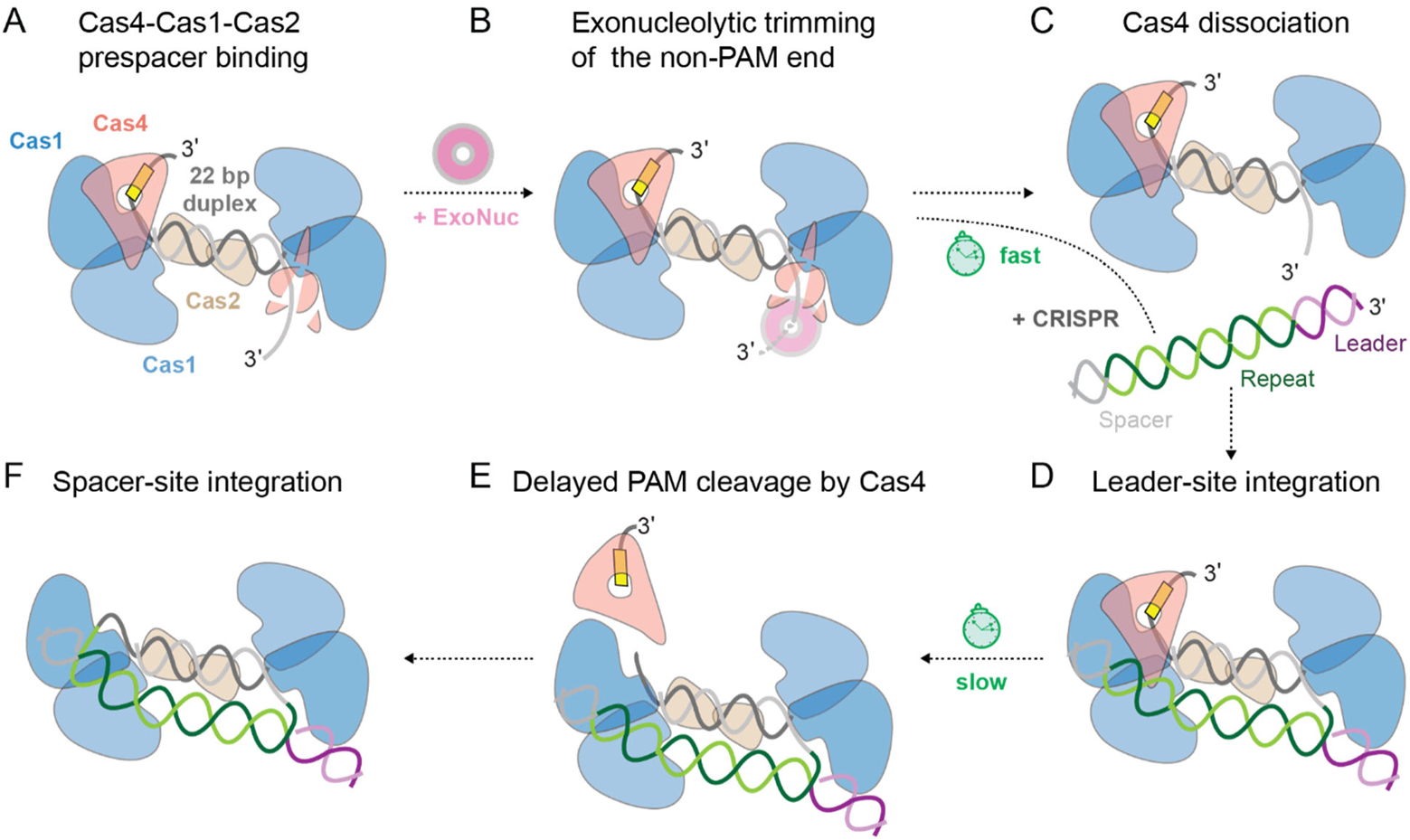
Kinetic model of Cas4-Cas1-Cas2 mediated processing and integration of prespacers. (A) Cas4-Cas1-Cas2 binding to a PAM/NoPAM prespacer. (B) Rapid trimming of non-PAM end by exonucleases, with Cas4 providing partial protection. (C) Dissociation of Cas4 from the non-PAM end following trimming. (D) Association of Cas4-Cas1-Cas2 to the CRISPR array with integration of a mature non-PAM end at the leader-repeat junction. (E) Cleavage of PAM by Cas4 and dissociation of Cas4, handoff of mature PAM end to Cas1 for integration. (F) Integration of the mature PAM end at repeat-spacer junction.

We demonstrate that interactions between the C-terminal helix of Cas4 and the inactive Cas1 subunit combined with interactions between Cas4 α4 and Cas2 α2 are necessary for efficient cleavage. The Cas4 C-terminal helix extends from the iron-sulfur cluster domain, which contains several of the PAM-recognition residues. Thus, docking of Cas4 to Cas1 via the C-terminal helix may position the iron-sulfur cluster domain correctly for PAM recognition. The position of Cas4 α4 is also important to allow ssDNA to pass through the Cas4 active site. Our cryo-EM reconstruction of the PAM/NoPAM complex suggests that this helix is conformationally flexible in the absence of PAM binding. Notably, the α4 and C-terminal helices are the most mobile structural elements in the five highest ranked AlphaFold models (**Figure S3A**). We propose that interactions formed by the Cas4 C-terminal and α4 helices with Cas1-Cas2 allow passage of the PAM through the Cas4 active site and stable recognition via a properly positioned iron-sulfur domain.

Our results indicate that Cas4 enhances prespacer integration by Cas1-Cas2 at the leader site. In native type I-A and I-B systems, deletion of Cas4 substantially reduces spacer acquisition (Li et al., 2014; Shiimori et al., 2018), although this reduction is less pronounced in a heterologous type I-D system (Kieper et al., 2018), potentially due to overexpression of Cas1-Cas2. Combined with our *in vitro* observations, these results suggest that Cas4 is necessary not only for prespacer selection and processing, but also for optimal integration of spacers in the CRISPR array. The observed enhancement is likely due to higher affinity of prespacer binding in the presence of Cas4 due to specific PAM interactions. Importantly, enhanced integration relies on delayed Cas4 cleavage activity, as Cas4 does not enhance integration of a fully processed prespacer. Thus, the slow kinetics of Cas4 cleavage enables asymmetric prespacer processing and enhanced leader site integration of the non-PAM end, contributing to the directional integration of the prespacer.

Recent structural and biochemical studies of the type I-G Cas4/1-Cas2 complex indicated that interactions between Cas4/1 and the CRISPR activates the Cas4 domain following half-site integration (Hu et al., 2021). However, we have not observed activation of type I-C Cas4 under many different conditions, suggesting that type I-G and I-C systems use alternative mechanisms for Cas4 activation. Importantly, we do not observe efficient cleavage activity of Cas4 at reduced temperatures (e.g. 37 °C), either for prespacer or HSI substrates. This suggests the possibility of another factor that might activate type I-C Cas4 for PAM cleavage at mesophilic temperatures. In type I-E and I-F systems, host factor IHF binds and bends the CRISPR leader DNA to enhance Cas1-Cas2 binding and allow efficient integration to occur (Fagerlund et al., 2017; Nuñez et al., 2016; Wright et al., 2017; Yoganand et al., 2017). It remains unknown if systems that contain additional factors like Cas4 require such host factors. However, various conserved upstream sequence elements have been identified in specific CRISPR-Cas subtypes, including type I-C, and may be important for spacer acquisition as observed in type I-E and I-F (Santiago-Frangos et al., 2021). Further investigation is required to determine whether Cas4-Cas1-Cas2 or a host factor interacts with upstream sequence elements in the CRISPR leader to activate Cas4 and ensure integration fidelity.

Delayed PAM processing is a common feature in type I systems to achieve directional spacer integration, although the exact mechanisms by which PAM processing is controlled remains unclear. In type I-E, which lacks Cas4, protection of the PAM end by the Cas1 C-terminal tail leads to delayed exonucleolytic trimming of the PAM end, ensuring integration of the non-PAM end at the leader site (Kim et al., 2020; Ramachandran et al., 2020). However, it is not known how the PAM is released from Cas1 and processed prior to the second integration event. Similarly it remains unclear how two distinct Cas4 proteins control prespacer directionality in type I-A (Shiimori et al., 2018), how the Cas4 domain in type I-G is activated following leader site integration (Hu et al., 2021), or whether an additional factor is required to activate Cas4 in type I-C. Future studies will be necessary to fully elucidate these mechanisms.

## Methods

### Cloning, protein expression and purification

Previously described constructs for Cas1, Cas2 and Cas4 expression were used to express and purify the individual proteins (Lee et al., 2018, 2019). For co-expression with Cas4, sufABCDSE genes were amplified from pSUF plasmid and cloned into pACYC using Gibson assembly. All primers used for cloning various constructs have been listed in Table S1.

Cas1 and Cas2 were overexpressed in *E. coli* BL21(DE3) and grown at 37 °C to 0.5 OD600 in LB media, followed by overnight induction at 17 °C with 1 mM IPTG. Cas4 was co-expressed with sufABCDSE-pACYC in *E. coli* C41 cells and grown to an OD600 of 0.5 in 2XTY media. Cells were induced with 1 mM IPTG and the cultures were supplemented with 100 mg of ferric citrate, ferrous sulfate, ferrous ammonium sulfate and L-cysteine at the time of induction. 1000x solutions for ferrous sulfate, ferrous ammonium sulfate and a 100x solution for ferric citrate were prepared and filtered prior to induction. Cultures were grown overnight at 17 °C. For all protein purifications, cells were harvested by centrifuging at 4000 rpm for 30 minutes, resuspended in Ni-NTA buffer (50 mM Na_2_HPO_4_ pH 8.0, 500 mM NaCl, 5% glycerol, 2 mM DTT) supplemented with 10 mM imidazole (pH 8.0) and 100 mM PMSF, and lysed using a sonicator (Branson Ultrasonics™ Sonifier™). The lysate was centrifuged at 18000 rpm for 30 minutes (Beckman Coulter Avanti J-E) and the supernatant was used for further purification. All proteins were initially purified using HisPur Ni-NTA affinity resin (Thermo Fisher Scientific) using Ni-NTA buffer supplemented with 25 mM imidazole during wash and 250 mM imidazole during elution. His_6_-MBP-Cas1 and His_6_-MBP-Cas2 were dialyzed against Ni-NTA buffer and His_6_-MBP was cleaved using TEV protease overnight at 4 °C. The cleaved protein was purified with subtractive Ni-NTA. Flow through and wash fractions were analyzed by SDS-PAGE. The cleanest fractions were pooled. In case of MBP contamination following subtractive Ni-NTA, the protein was dialyzed in buffer A (20 mM HEPES pH 7.5, 100 mM KCl, 5% glycerol and 2 mM DTT) for 3 hours at 4 °C. Dialyzed protein was loaded on a 5 mL HiTrap Heparin column (GE Healthcare) equilibrated with buffer A to remove the MBP, and Cas1 or Cas2 was eluted using a gradient of buffer B (20 mM HEPES pH7.5, 2M KCl, 5% glycerol, 2 mM DTT). The protein was concentrated and further purified on a Superdex 75 16/600 GL column (GE Healthcare), in a size exclusion buffer (20 mM HEPES pH 7.5, 100 mM KCl, 5% glycerol and 2 mM DTT).

To reduce the possibility of protein purification tags on Cas1 or Cas4 interfering with activation of Cas4 in the presence of a half site intermediate, we redesigned our constructs to His_6_-SUMO-Cas1 and Cas4 with His_6_ on the C-terminus. For His_6_-SUMO-Cas1, Ni-NTA purified protein was dialyzed overnight against Ni-NTA buffer and the SUMO tag was cleaved using Ulp1 protease. The cleaved protein was purified with subtractive Ni-NTA followed by purification on a Superdex 75 16/600 GL column (GE Healthcare), in a size exclusion buffer (20 mM HEPES pH 7.5, 100 mM KCl, 5% glycerol and 2 mM DTT). For His_6_-Cas4 with His tag on N- or C-terminus, NiNTA purification was directly followed by further purification on Superdex 200 16/600 GL column (GE Healthcare), in a higher salt size exclusion buffer (20mM HEPES pH 7.5, 500 mM KCl, 5% glycerol and 2 mM DTT).

ExoT was obtained from New England Biolabs.

### Complex formation

For Cas4-Cas1-Cas2 complex formation, DNA substrates were pre-annealed and purified using native 10% PAGE. For assembling complexes with various DNA substrates, 20 µM Cas1, 10 µM Cas2, 30 µM Cas4, and 10 µM substrate DNA were mixed (final approximate ratio of 2:1:3:1) in a final volume of 500 µL in complex formation buffer (20 mM HEPES (pH 7.5), 100 mM KCl, 5% glycerol, 2 mM DTT, and 2 mM MnCl_2_) and incubated on ice for 45 min. Cas4-Cas1-Cas2-DNA complexes were purified using a Superdex 75 10/300 GL column (GE Healthcare) in complex formation buffer. Peak fractions were visualized by SDS-PAGE and the fractions containing all three proteins were pooled and concentrated to 8-10 µM, flash frozen in liquid nitrogen and stored at −80°C.

### Cryo EM grid preparation and data acquisition

3 µl of ~ 1.5 µM (0.75 mg/ml) of the complex was applied to freshly glow discharged quantifoil 300 mesh R 1.2/1.3 grids (EMS) and plunge frozen into liquid ethane using a Thermo Fisher Vitrobot Mark IV with conditions of 4 °C, 100% humidity, blot force 3, and a 3 second blot time. Grids were clipped and transferred to Thermo Fisher Scientific 200 kV Glacios microscope equipped with a K3 Direct Electron Detector (DED) at the Iowa State University cryo-EM facility. 45 frame movies were recorded in super-resolution counting mode with pixel size of 0.44865 Å. Total electron dose was 50.5 e^−^ Å^2^. Movies were collected using SerialEM (Mastronarde, 2005) with single image per hole strategy and a defocus range of 1.5-3.5 µm.

### Cryo-EM data processing

All data processing was performed in cryoSPARC v3.1.0 and v3.2.0 (Punjani et al., 2017). Following motion correction and CTF estimation, blob picking was performed with a circular blob of minimum and maximum particle diameter of 100 Å and 300 Å respectively for the first data set. Particles were then subjected to multiple rounds of 2D classifications with a 200 Å circular mask diameter and resulting classes were used for template picking with a particle diameter of 300 Å. Template picked particles were extracted with a box size of 384 and binned to a box size of 128 for processing until homogenous refinements. Extracted particles were subjected to iterative 2D classifications. Selected 2D classes were then used to build an ab initio model. All particles were refined using homogenous refinement with the ab initio model. These were then used for heterogeneous refinements into 3-5 classes depending on the number of particles in the dataset. Homogenous and Non-uniform refinements (Punjani, Zhang and Fleet, 2020) were further used with CTF refinement of particles performed before the refinement step or on the fly during the refinement. Figures S2, S5, S6, S7 and S9 illustrate the processing pipeline for PAM/PAM, PAM/PAM with 24 bp duplex, PAM/NoPAM, PAM/Proc and HSI maps respectively. Data collection details and validation statistics for all reconstructions have been listed in Table 1. All maps were visualized using UCSF Chimera during processing (Pettersen et al., 2004). Figures were prepared in Chimera X (Goddard et al., 2018).

**Table 1:** Data collection parameters for the five reconstructions and refinement statistics for four models described in this study

|  | PAM/PAM | PAM/NoPAM | PAM/Proc | HSI | PAM/PAM<br>24bp duplex |
| --- | --- | --- | --- | --- | --- |
| **Data collection and processing** |  |  |  |  |  |
| Magnification | 45000 | 45000 | 45000 | 45000 | 45000 |
| Voltage (kV) | 200 | 200 | 200 | 200 | 200 |
| Electron exposure (e-/Å<sup>2</sup>) | 50 | 50 | 50 | 50 | 50 |
| Defocus range | ~1.5 to 3.5 | ~1.5 to 3.5 | ~1.5 to 3.5 | ~1.5 to 3.5 | ~1.5 to 3.5 |
| Pixel size (Å) | 0.9063 | 0.9063 | 0.9063 | 0.9063 | 0.9063 |
| Symmetry imposed | C1 | C1 | C1 | C1 | C1 |
| No of movies collected | 1571 | 1506 | 1554 | 1947 | 6960 |
| Final map resolution (Å) | 3.46 | 3.9 | 3.41 | 4.3 | 3.9 |
| FSC threshold | 0.143 | 0.143 | 0.143 | 0.143 | 0.143 |
| **Refinement** |  |  |  |  |  |
| B factors (Å<sup>2</sup>) |  |  |  |  |  |
| Protein (max) | 95.05 | 178.17 | 142.90 | 185.76 | N/A |
| Protein (mean) | 24.22 | 96.65 | 73.98 | 95.89 |  |
| RMS Deviations |  |  |  |  |  |
| Bond lengths (Å) | 0.004 | 0.006 | 0.006 | 0.007 | N/A |
| Bond angles (°) | 0.91 | 0.72 | 1.00 | 1.14 | N/A |
| Validations |  |  |  |  |  |
| MolProbity | 1.87 | 2.13 | 2.06 | 3.20 | N/A |
| Clashscore | 8.8 | 14.21 | 11.11 | 14.83 | N/A |
| Poor rotamers (%) | 0.06 | 0.00 | 0 | 2.7 | N/A |
| Model Composition |  |  |  |  |  |
| Non-hydrogen atoms | 17485 | 17153 | 15518 | 17375 | N/A |
| Amino acid residues | 2001 | 1974 | 1784 | 1790 | N/A |
| Ligands | 4 | 3 | 2 | 3 | N/A |
| Nucleotides | 65 | 60 | 56 | 146 | N/A |
| Ramachandran plot |  |  |  |  |  |
| Favored (%) | 94.06 | 92.43 | 91.75 | 87.44 | N/A |
| Allowed (%) | 5.74 | 7.36 | 7.97 | 10.53 | N/A |
| Disallowed (%) | 0.2 | 0.20 | 0.28 | 2.03 | N/A |

### Modeling, Refinement, and analysis

AlphaFold (Jumper et al., 2021; Mirdita et al., 2022; Varadi et al., 2022) models for Cas1 and Cas4 and the crystal structure for Cas2 (Nam et al., 2012) were used as templates to facilitate building models for the three proteins in *Coot* (Emsley et al., 2010). DNA for the prespacer (PAM/PAM, PAM/NoPAM, PAM/Proc models) and the CRISPR array (HSI model) were built de novo in Coot. Multiple rounds of model refinement were performed in Phenix.real_space_refine (Liebschner et al., 2019) with secondary structure, base pairs, stacking pairs and Fe_4_S_4_ cluster restrains. Molprobity (Williams et al., 2018) was used to guide iterative refinements in Phenix with manual adjustments performed in Coot between subsequent rounds of refinements. Adjustments to the DNA sequence and removal of bases or protein chains for specific models were done in Coot followed by refinements in Phenix. All figures with density maps or models or both were prepared using ChimeraX (Goddard et al., 2018).

### DNA substrate preparation

All oligonucleotides were synthesized by Integrated DNA Technologies. Sequences of all DNA substrates are shown in Table 2. All DNA substrates were first purified using 10% PAGE with 8 M urea and 1X TBE and ethanol precipitation. Purified prespacers were labeled with [γ-^32^P]-ATP (PerkinElmer) and T4 polynucleotide kinase (NEB) at the 5′ end. Excess ATP was removed using Microspin G-25 columns (GE Healthcare).

### Prespacer processing assays

For preparing duplex substrates, a 5ʹ radiolabeled strand was annealed an unlabeled complementary strand in a 1:2 ratio (25 nM radiolabeled strand and 50 nM unlabeled strand) in cleavage buffer (20 mM HEPES (pH 7.5), 100 mM KCl, 5% glycerol, 4 mM DTT, 4 mM MnCl_2_,). For processing assays with the half-site intermediate, the substrate was prepared by annealing the labelled strand with twice as much of each of the unlabeled DNA strands. The hybridization reactions were incubated at 95 °C for 3 min followed by slow cooling to room temperature.

Cas1, Cas2 and Cas4 were premixed on ice, substrate was added at a final concentration of 1-5 nM and the reactions were incubated at 65 °C for 30 min unless indicated otherwise. A final concentration of 250 nM Cas1, 250 nM Cas2 and 500 nM Cas4 was used unless otherwise indicated. Reactions were quenched with 2X RNA loading dye (95% formamide, 0.01% SDS, 0.01% bromophenol blue, 0.01% xylene cyanol supplemented with 25 mM of EDTA) and heated at 95 °C for 5 min. Samples were run on 10% urea-PAGE. The gels were dried and imaged using phosphor screens on a Typhoon imager.

For quantification, the intensity of bands was measured by densitometry using ImageJ (Schneider, Rasband and Eliceiri, 2012). The fraction cleaved was calculated by dividing the product band by the sum of both bands. The values from three or four replicates were averaged, error is reported as standard deviation between the replicates.

For comparison of PAM/NoPAM, PAM/Proc and HSI substrates (Figure 6D), the reactions were performed at a variety of temperatures and Cas1, Cas2 and Cas4 concentrations. We did not observe differences in activity regardless of protein concentration, and we did not observe processing activity at mesophilic temperatures for any substrate tested. Concentrations of 100 nM Cas1, 100 nM Cas2 and 50 nM Cas4 at a temperature of 65 °C were used for the experiment shown in Figure 6D. Cas1 purified using both His_6_-MBP-Cas1, which retains additional GAGS N-terminal amino acids following TEV cleavage, and His_6_-SUMO-Cas1 constructs, which does not contain any additional sequence on the N- or C-terminus, were tested and the assays done with His_6_-SUMO-Cas1 are reported in Figure 6D-E. Cas4 purified with both N- and C-terminal His tags were tested and the cleavage assays reported in Figure 6D-E were done with C-terminal His tag.

All oligonucleotides used for the assays are listed in Table S2.

### Exonuclease trimming assays and integration assays using PAM/Proc substrates

Reactions were performed in buffer containing 20 mM HEPES (pH 7.5), 100 mM KCl, 5% glycerol, 10 mM MgCl_2_, 5 mM MnCl_2_ and 2 mM DTT. Purified Cas1 (100 nM), Cas2 (50 nM), and Cas4 (50 nM) were pre-incubated with 5 nM 5′-radiolabelled prespacer DNA on ice for 15 min. For the exonuclease trimming assays, ExoT was added at a concentration of 1 U/µL, and the reactions were further incubated at 37 °C for 5 min, 15 min and 30 min. To observe integration, 1 µM of the linear mini-CRISPR array (consisting of a 20 bp leader segment, a 32 bp repeat sequence and a 5 bp spacer sequence, shown in Table S2) was added to the reaction prior to incubation at 37°C. For the exonuclease trimming assays, the mini-CRISPR DNA contained phosphorothioate groups at the 3′ ends to prevent degradation by the exonuclease. For the time course reactions, a master mix was prepared and an aliquot of 10 µL was taken at each time point. Reactions were quenched with 2X loading dye at the indicated time points. The samples were denatured at 95 °C for 5 minutes and analyzed by 10% urea-PAGE, as described above. Oligonucleotides used for the assays have been listed in Table S2.

## Acknowledgements

We thank Dr. Eric Underbakke and the lab for the generous gift of pET-His-SUMO vector and Ulp1 protease. We thank Dr. Yang Yang for helpful discussions on model building with cryo-EM reconstructions; Dr. Scott Nelson and Dr. Baoyu (Stone) Chen for helpful discussions on experimental design. We thank members of the ResearchIT group at Iowa State University for computational support; Tracey Stewart and Roy J Carver High Resolution Microscopy Facility for technical assistance with negative stain EM screening of samples used in cryo-EM. We thank all the members of the Sashital Lab for helpful input throughout the course of this project. The Iowa State University Cryo-EM Facility was established through a grant from the Roy J. Carver Charitable Trust (Muscatine, IA). This work was supported by an NIH R01 grant (GM115874) and NIH R35 grant (GM140876) to DGS.

## Author Contributions

Y.D. performed protein purification, complex reconstitution, and grid preparation. Y.D. and P.J. performed grid screening for cryo-EM data collection. P.J. collected the cryo-EM data. Y.D. performed the cryo-EM data processing with P.J. and D.G.S. contributing. D.G.S. built the atomic models. D.G.S and Y.D. refined the models. Y.D. performed all the cleavage assays. S.K.S. performed exonuclease trimming and integration assays. Y.D., S.K.S. and D.G.S analyzed and interpreted the biochemical results, wrote, and edited the manuscript. D.G.S supervised the research and secured funding.

## Declaration of interests

The authors declare no competing interests.

## Notes

### Competing Interest Statement

The authors have declared no competing interest.

