## Supplemental Information for "PAM binding ensures orientational integration during Cas4-Cas1-Cas2 mediated CRISPR adaptation"

### Supplementary Figures

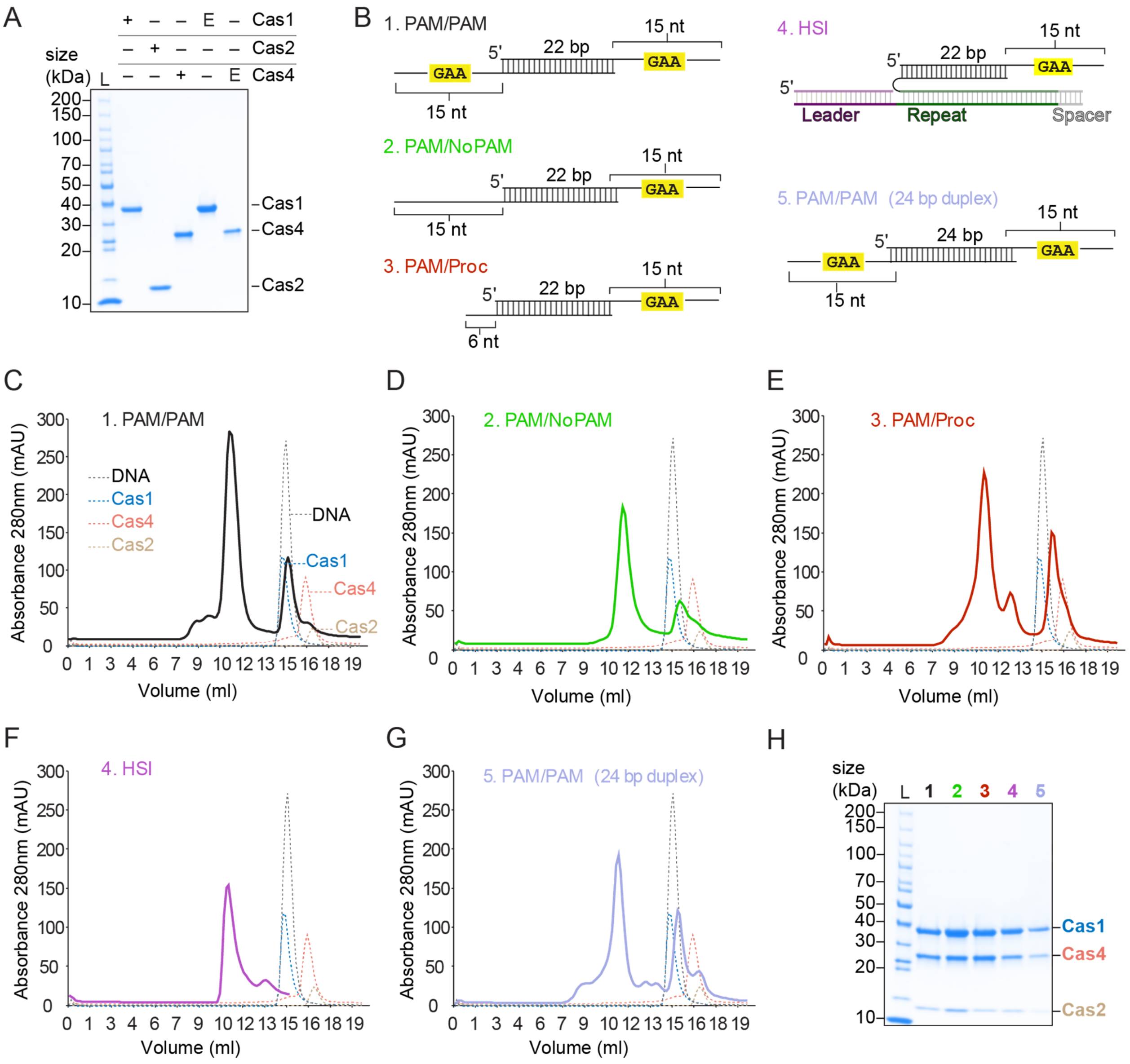


**Figure S1, related to Figure 1, 4, 5 and 6: Cas4-Cas1-Cas2 complex formation with various substrates** (A) Coomassie blue stained SDS polyacrylamide gel for purified Cas1, Cas2, Cas4 and active site mutants of Cas1 (E166A) and Cas4 (E108A). (B) Schematic of substrates used in this study for cryo-EM reconstructions. (C-G) Size exclusion chromatography (SEC) of various complexes shown in this study, overlay with SEC of individual components. SEC was performed on Superdex 200 Increase 10/300 GL. (H) Coomassie blue stained SDS polyacrylamide gel for samples used for grid preparation for cryo-EM.

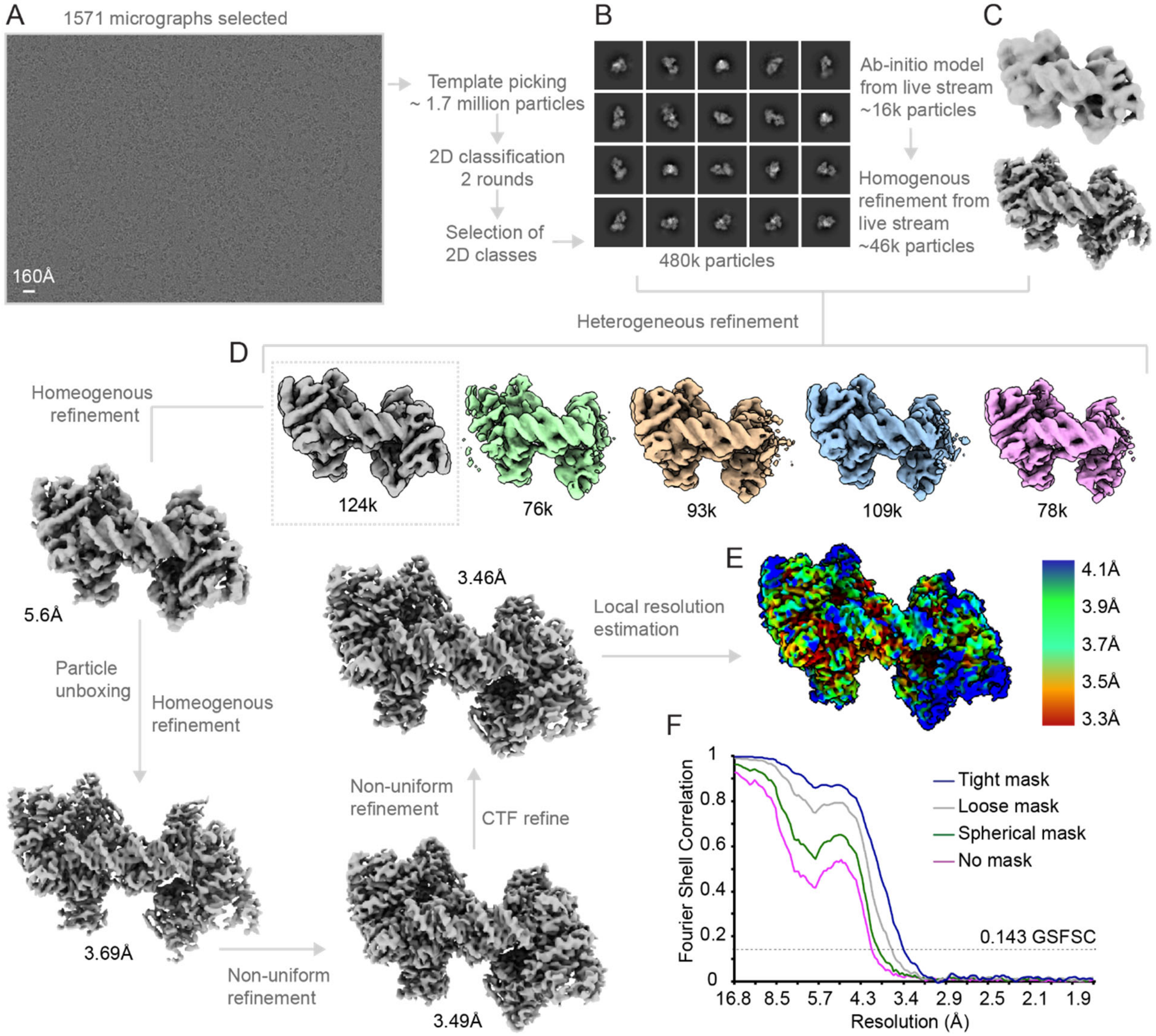


**Figure S2, related to Figure 1: Cryo-EM data processing of PAM/PAM complex** (A) Representative micrograph. (B) Selected 2D classes with a total of 480k particles used for further processing. (C) Ab initio models generated during live stream processing of the dataset. (D) Heterogeneous refinement of selected particles in (B) with ab initio model in (C), 3D class with 124k particles selected and used for further homogenous refinements, particle unboxing and non-uniform refinement (with CTF refinement on the fly). (E) Local resolution estimation of final map. (F) FSC values plotted against resolution for final map with and without masks.

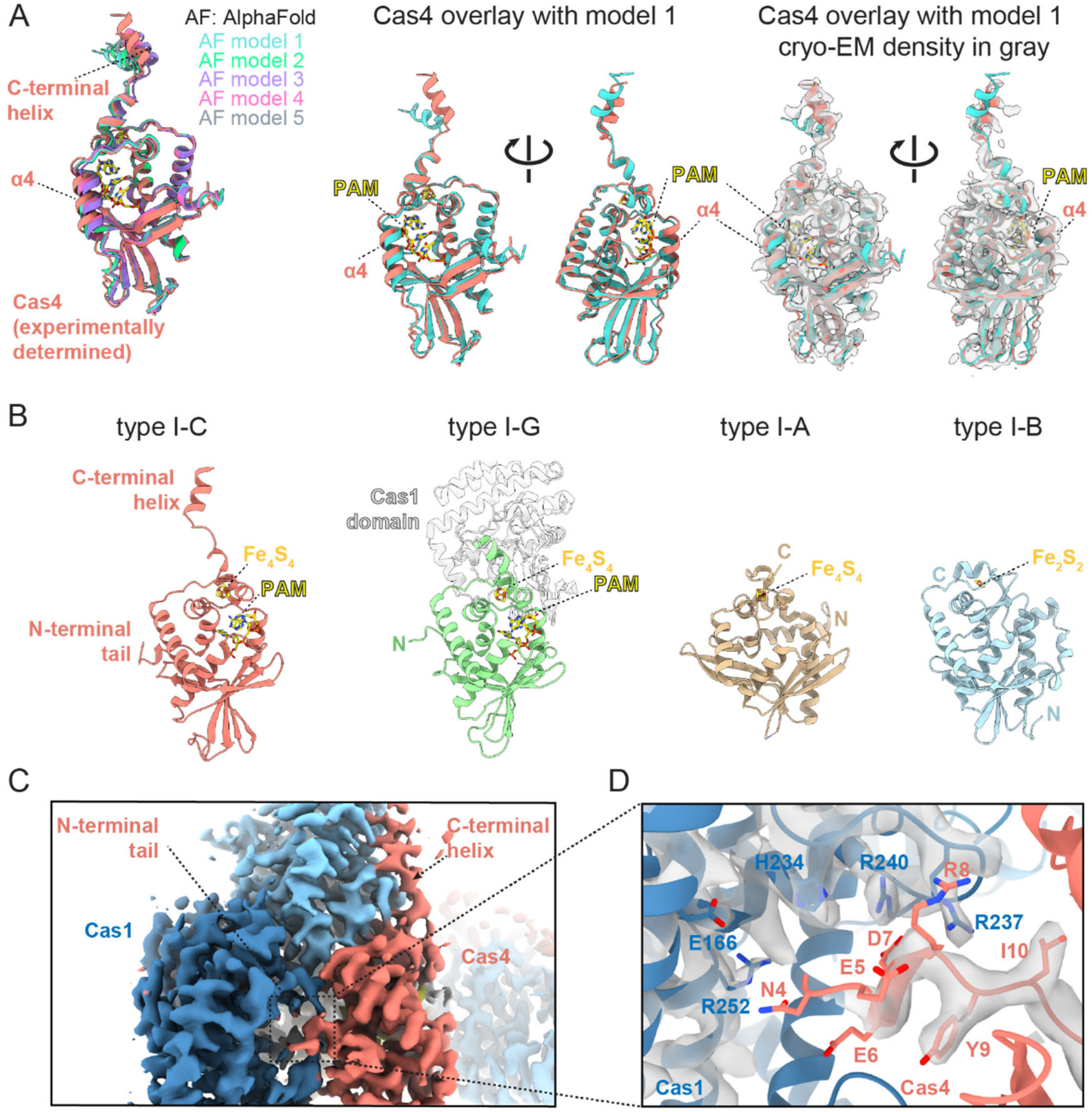


**Figure S3, related to Figure 1: Structural features of the type I-C Cas4 structure.** (A) Overlay of Cas4 structural model determined using the cryo-EM reconstruction (salmon) and AlphaFold predicted models. Front and back views of overlay with highest ranking AlphaFold model are shown with and without the cryo-EM density. (B) Comparison of known Cas4 structures from type I-C (this study), type I-G (Hu *et al.*, 2021, PDB: 7MI4), type I-A (Lemak *et al.*, 2013, PDB: 4IC1), and type I-B (Lemak *et al.*, 2014, 2014, PDB: 4R5Q). The type I-G Cas4 structure is a domain of a Cas4/1 fusion protein. The Cas1 domain is shown in light gray. The N-terminal tail and C-terminal helix are labeled for type I-C Cas4. N- and C-termini are labeled for other Cas4 structures.

Iron-sulfur clusters and PAM sequences (for type I-C and I-G) are shown as sticks and labeled. (C) Close-up of cryo-EM density for Cas4 interaction with Cas1. Density is shown for the Cas4-Cas1-Cas2 complex bound to the PAM/processed substrate, which had the best resolution for PAM-bound Cas4. (D) Cryo-EM density for the Cas4 N-terminal tail interaction with Cas1. Density is shown for the N-terminal tail of Cas4, and Cas1 helices and loops containing catalytic residues E166 and H234 and basic residues that contribute to the electropositive groove.

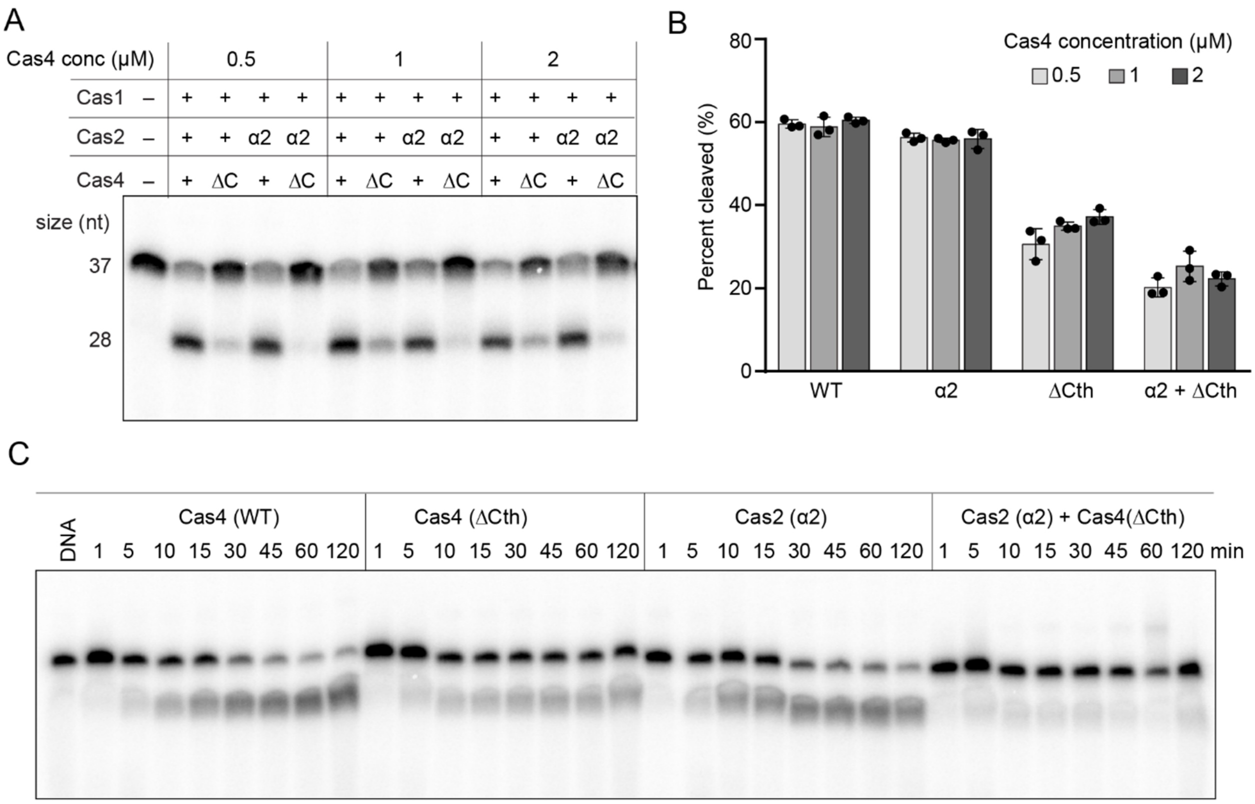


**Figure S4, related to Figure 2: Cas4 activation mediated by interactions with Cas1 and Cas2** (A) Polyacrylamide gel showing cleavage assay for individual Cas2 and Cas4 mutants and combination of the two at different concentrations of Cas4. ΔC indicates Cas4 C-terminal deletion. α2 indicates helix 2 mutant of Cas2. (B) Quantification of percent cleaved for assay shown in (A). Data is representative of three replicates. Error bars represent standard deviation and the three data points are plotted as dots. **(C)** Representative denaturing polyacrylamide gel showing cleavage assay for mutants at various time points. Quantification shown in Figure 2D.

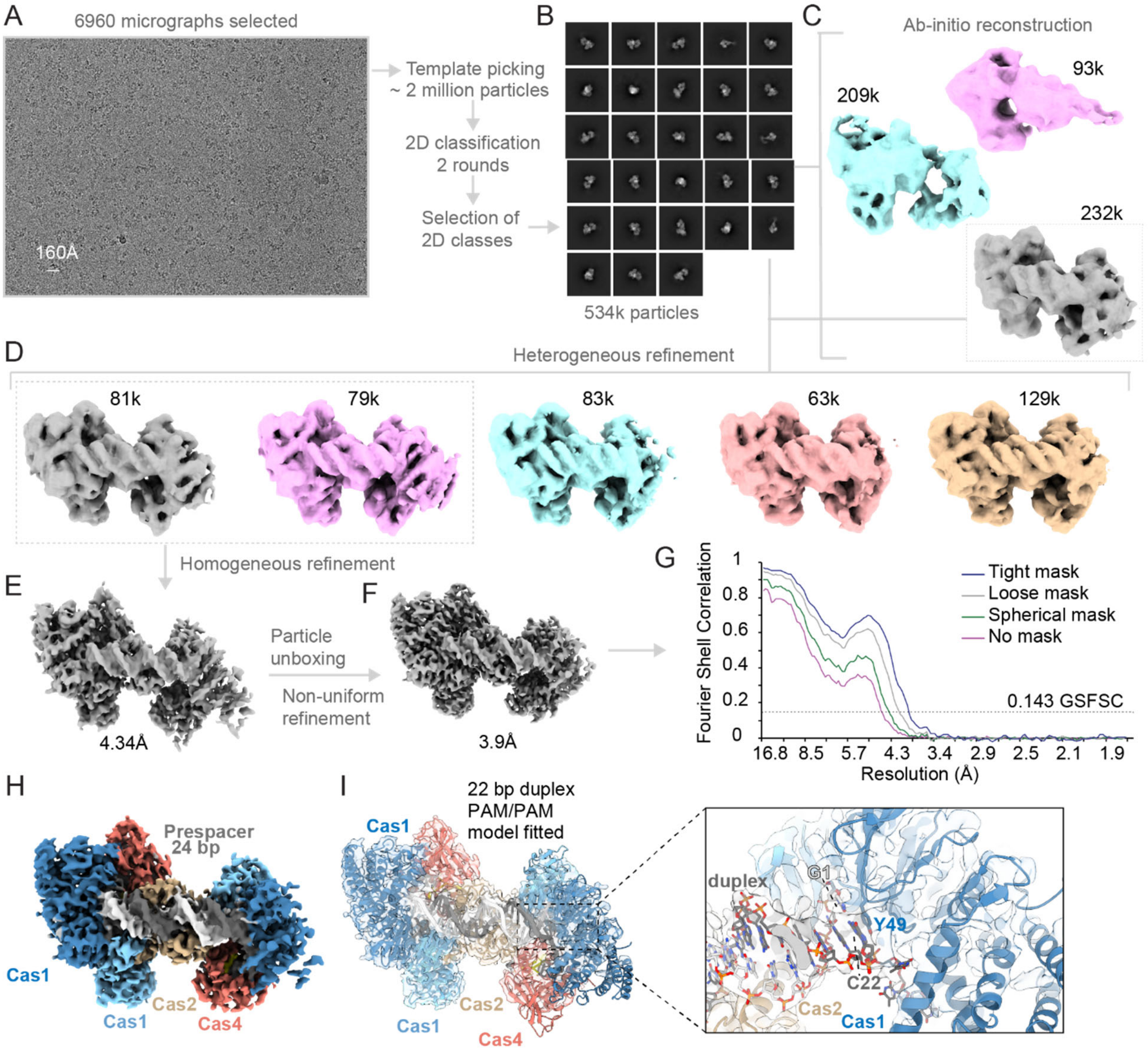


**Figure S5, related to Figure 4: Cryo-EM data processing of PAM/PAM 24 bp duplex complex**

(A) Representative micrograph. (B) Selected 2D classes with a total of 404k particles used for further processing. (C) Ab initio models generated using the selected 404k particles. (D) Heterogeneous refinement of selected particles in (B) with ab initio model in (C). (E) Homogeneous refinement of particles from selected 3D classes. (F) Non-uniform refinement, final map for PAM/PAM with 24 bp duplex reconstruction. (G) FSC values plotted against resolution for final map with and without masks. (H) Architecture of Cas4-Cas1-Cas2 within the 24 bp duplex reconstruction. (I) Fitted model for PAM/PAM 22 bp duplex into the 24 bp map, close up view of last base pair of a 22bp duplex stacking against Tyr49 of Cas1.

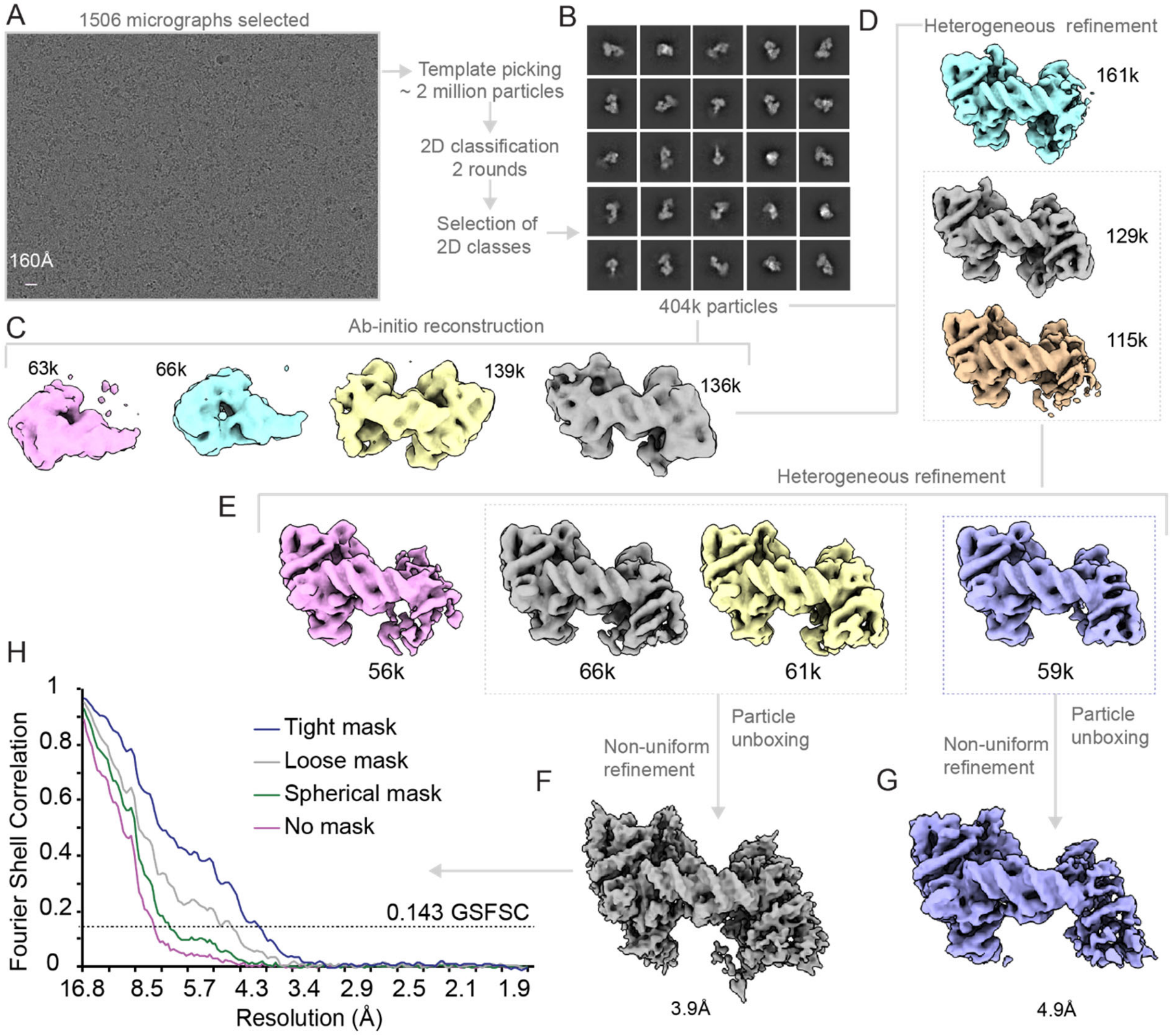


**Figure S6, related to Figure 5: Cryo-EM data processing of PAM/NoPAM complex** (A) Representative micrograph. (B) Selected 2D classes with a total of 404k particles used for further processing. (C) Ab initio models generated using the selected 404k particles. (D) Heterogeneous refinement of selected particles in (B) with ab initio model in (C). (E) Heterogeneous refinement of particles from selected 3D classes. (F) Non-uniform refinement of 3D classes with only partial density for Cas4 on one side, final map for PAM/NoPAM reconstruction. (G) Non-uniform refinement of 3D classes with no Cas4 density on one side. (H) FSC values plotted against resolution for final map with and without masks.

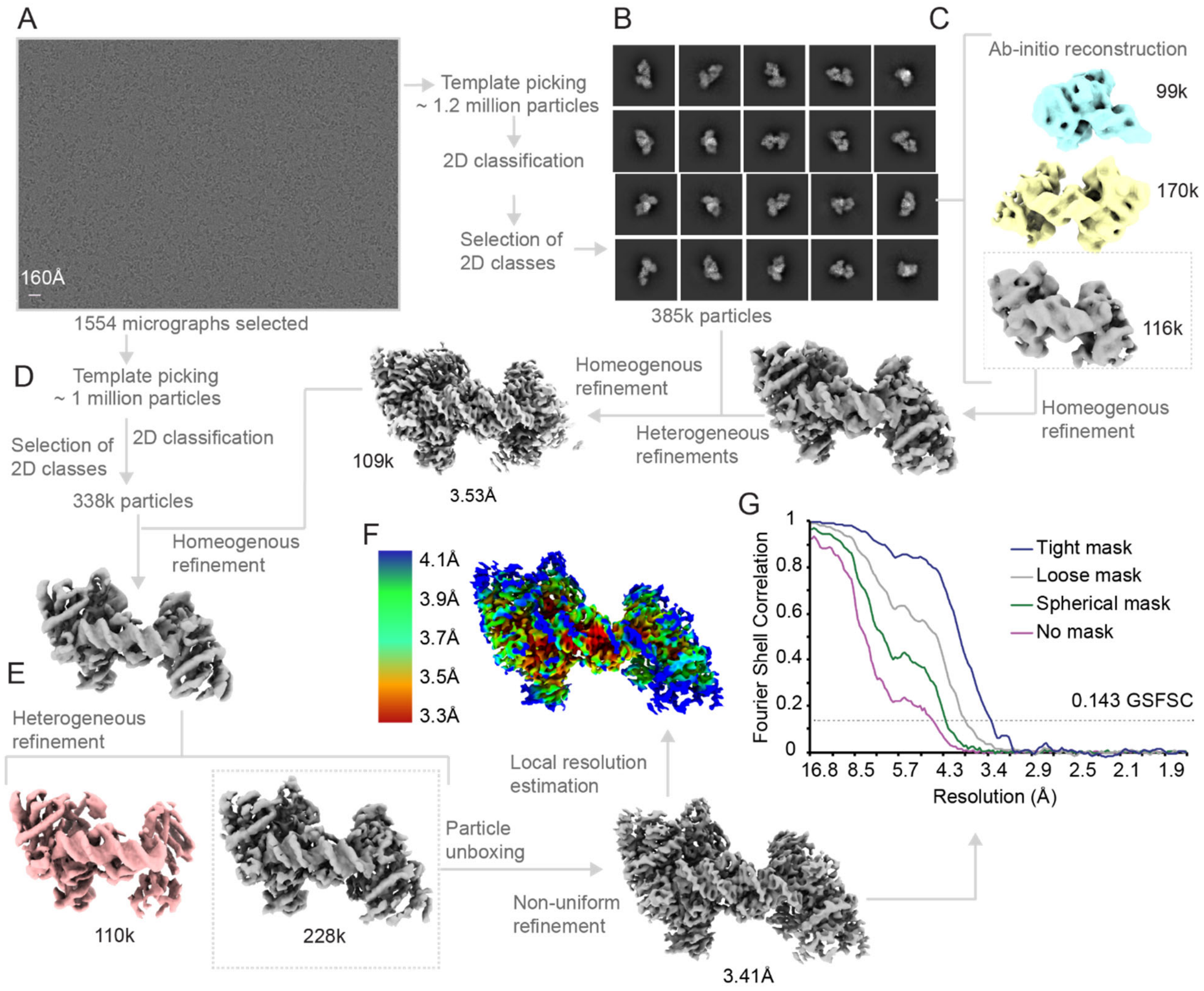


**Figure S7, related to Figure 5: Cryo-EM data processing of PAM/processed complex** (A) Representative micrograph. (B) Selected 2D classes with a total of 385k particles used for further processing. (C) Ab initio models generated using the selected 385k particles, selection of ab initio model for further heterogeneous refinements with selected particles in (B). (D) New round of template picking and selection of 2D classes, homogenous refinement of new particles with final map from (C). (E) Heterogeneous refinement of particles from (D), selection of class with strong density for all protein subunits and Cas4 density only just one side, non-uniform refinement of selected class, final map for PAM/processed reconstruction. (F) Local resolution estimation of final map. (G) FSC values plotted against resolution for final map with and without masks.

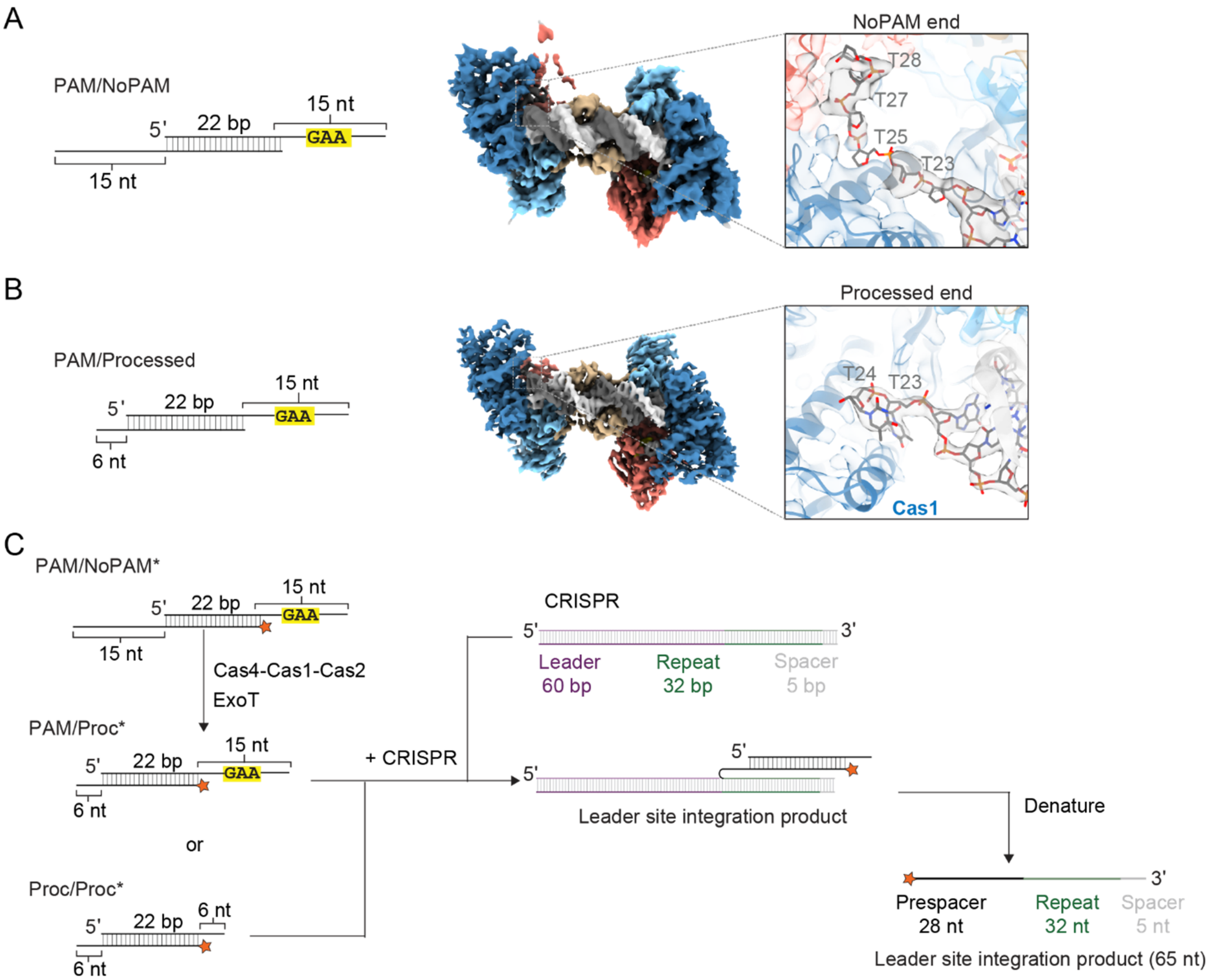


**Figure S8, related to Figure 5: Close up of non-PAM and processed ends and schematics for integration assays** (A) Substrate schematic and cryo-EM reconstruction of Cas4-Cas1-Cas2 PAM/NoPAM complex, zoom of density and model for non-PAM end of the PAM/NoPAM substrate within the reconstruction. The structure models the trajectory of the DNA backbone through the cryo-EM density. Nucleotides are labeled in gray. (B) Substrate schematic and cryo-EM reconstruction of Cas4-Cas1-Cas2 PAM/processed complex, zoom of density and model for processed end of the PAM/processed substrate within the reconstruction. The first two nucleotides of the 6 nt overhang are labeled. (C) Schematic of integration assay with PAM/NoPAM, PAM processed (PAM/Proc) or processed/processed (Proc/Proc) substrate. Asterisk indicates labeled strand. Radiolabel is indicated with a star in the substrate design.

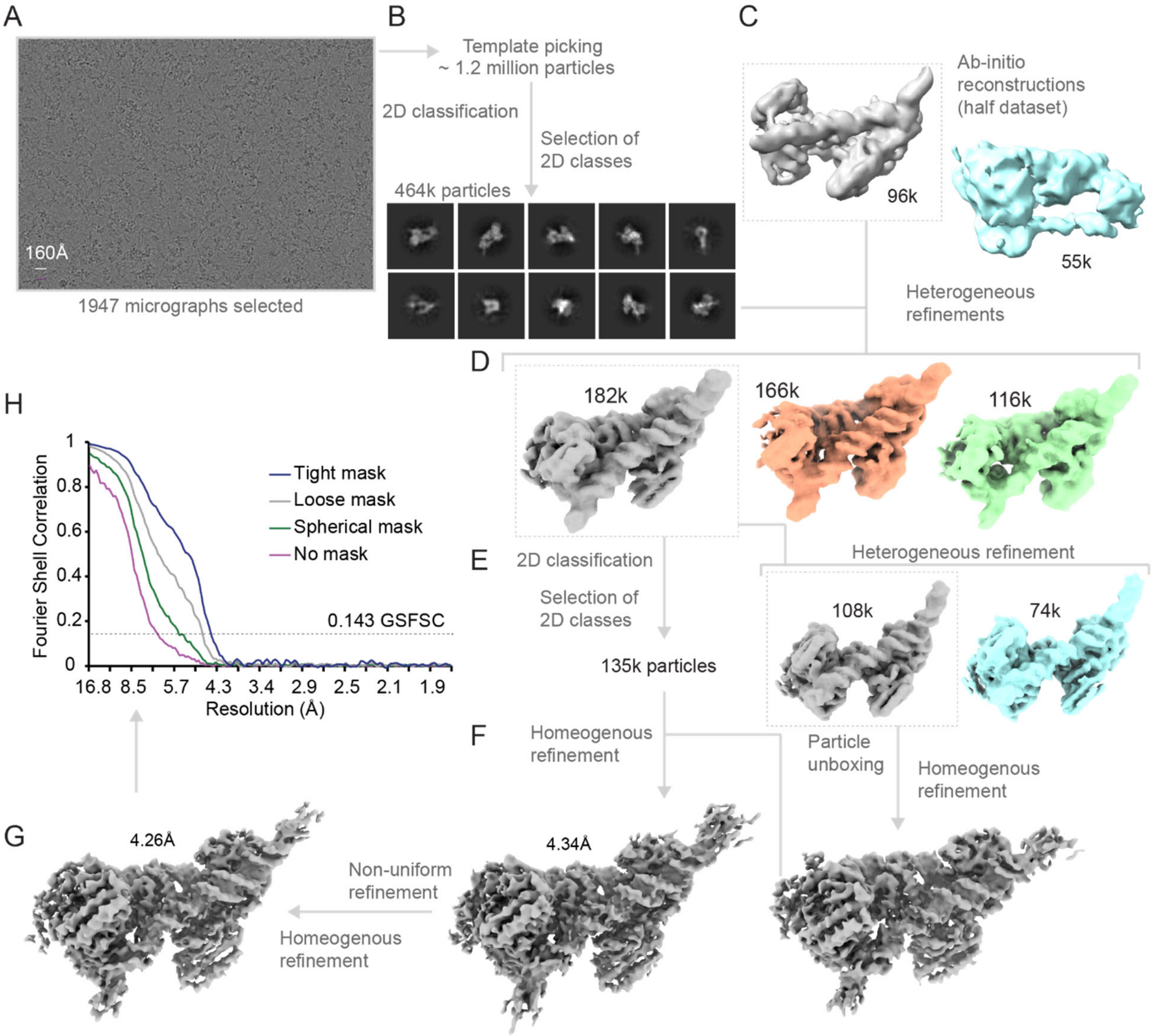


**Figure S9, related to Figure 6: Cryo-EM data processing of HSI complex** (A) Representative micrograph. (B) Selected 2D classes with a total of 464k particles used for further processing. (C) Ab initio models generated using the selected 464k particles, selection of ab initio model for further refinements. (D) Heterogeneous refinements of particles from (B). (E) 2D classification of particles in selected 3D class from (D), heterogeneous refinement of 135k selected particles of selected 2D classes. (F) Homogenous refinements for map obtained from heterogeneous refinement. (G) Non-uniform refinement and homogenous refinement for final map. (H) FSC values plotted against resolution for final map with and without masks.

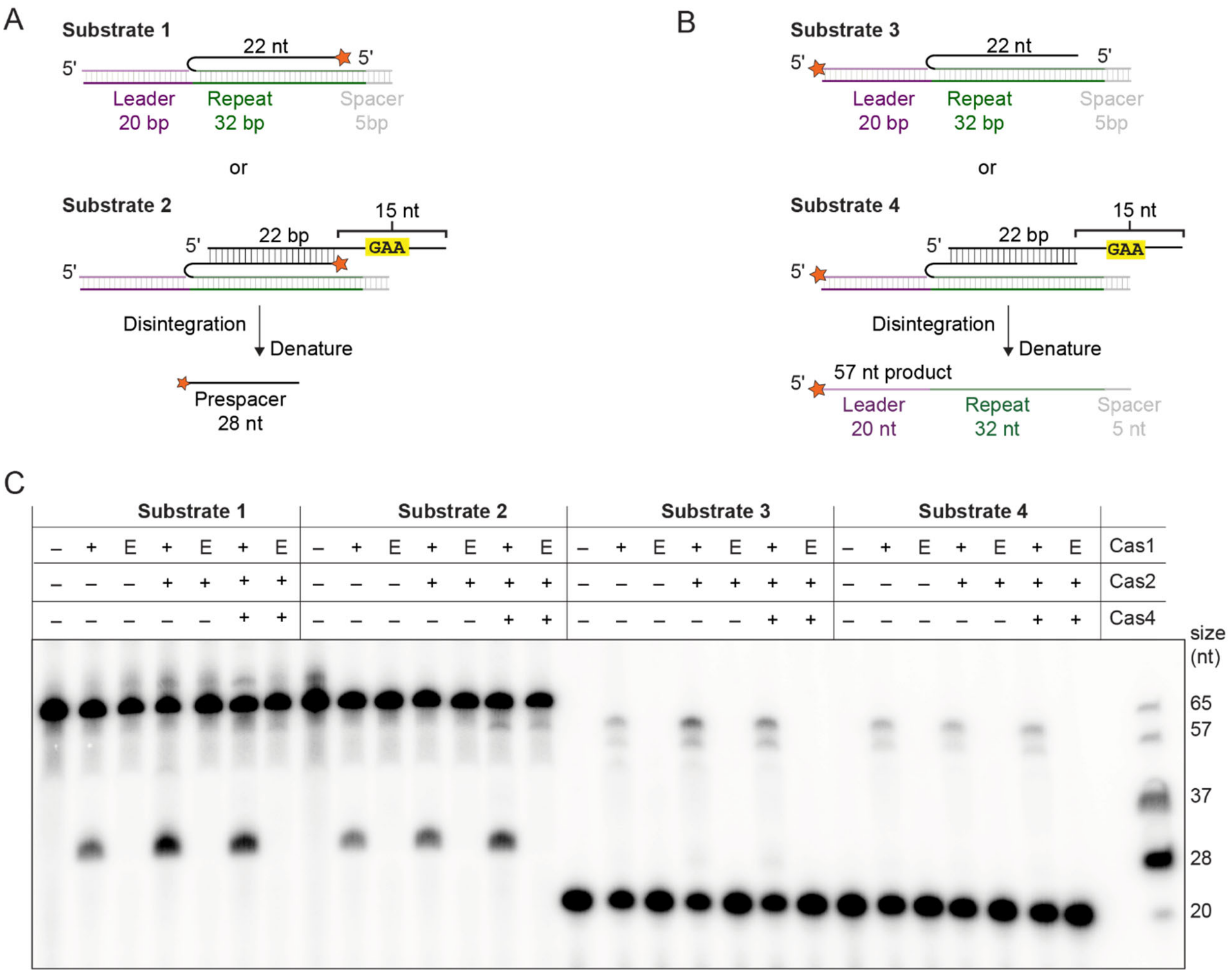


**Figure S10, related to Figure 6: Disintegration assays with and without Cas4** (A) Schematic of disintegration assay for substrate with radiolabel on prespacer strand. (B) Schematic for disintegration/integration assay with radiolabel on leader. (C) Polyacrylamide gel showing disintegration assay with Cas1 only; Cas1 and Cas2; or Cas1, Cas2 and Cas4. Cas1 active site mutant E166A is labeled E.

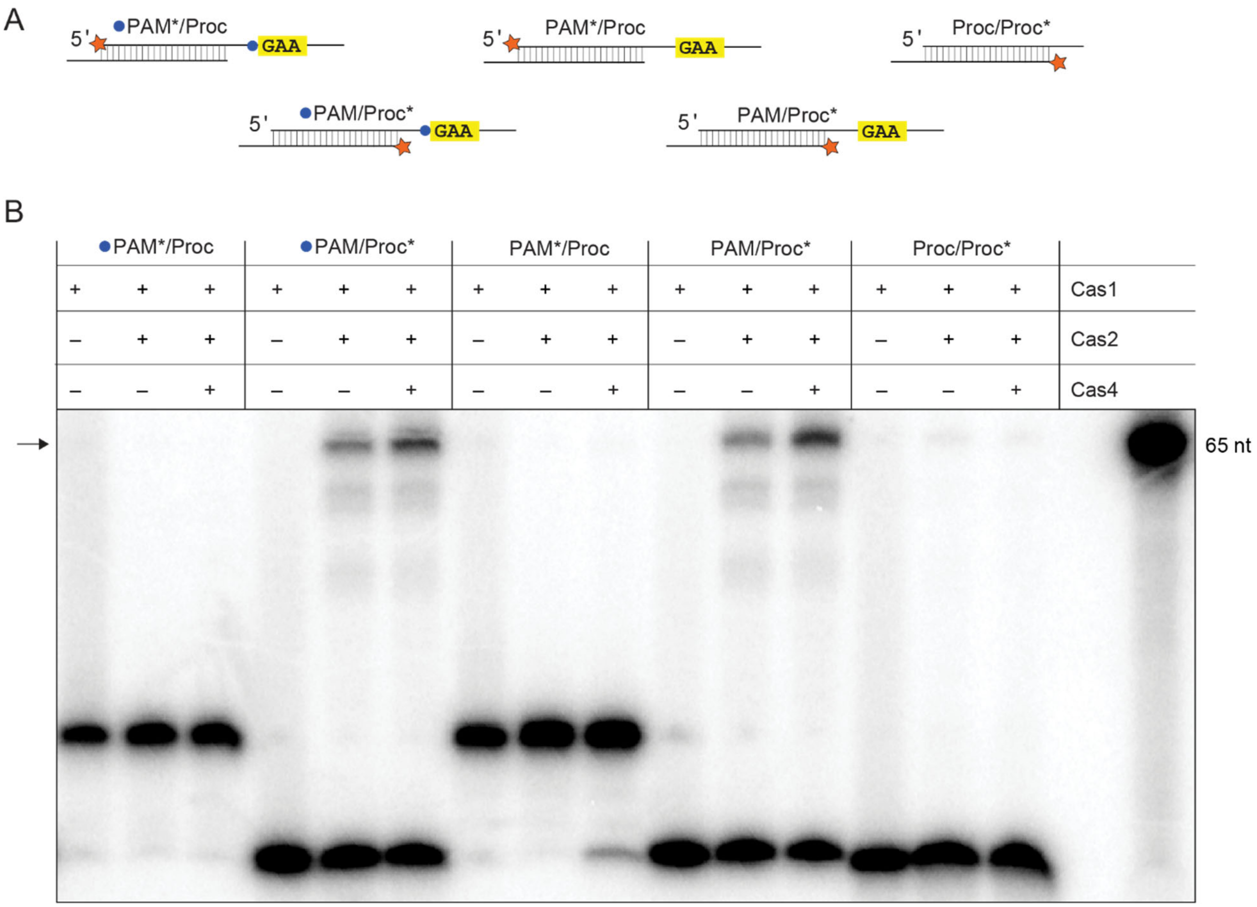


**Figure S11, related to Figure 6: Integration assay with blocked Cas4 cleavage activity** (A) Schematics of prespacer substrates tested for cleavage and integration. Phosphorothioate linkage is shown with a blue circle and radiolabel is indicated with an orange star. (B) Polyacrylamide denaturing gel showing integration of prespacer substrates in (A) in the absence and presence of Cas4. Black arrow indicates integration products.

**Table S1. Primers used in this study.**

| Name | Sequence (5′ → 3′) Description | Description |
|---|---|---|
| 1 | TAAGGAGATATACCATGGACATGCATTCAG GAACCTTTAACC | sufABCDSE forward for Gibson assembly pACYC/sufABCDSE |
| 2 | AGCAGCCTAGGTTAATTAATTAGCTAAGTGC AGCGGCTTTGG | sufABCDSE reverse for Gibson assembly pACYC/sufABCDSE |
| 3 | AGCCGCTGCACTTAGCTAATTAATTAACCTA GGCTGCTGCCACCG | pACYC forward for Gibson assembly pACYC/sufABCDSE |
| 4 | AATGCATGTCCATGGTATATCTCCTTATTAA AGTTAAACAAAATTATTTC | pACYC reverse for Gibson assembly pACYC/sufABCDSE |
| 5 | TTAGCAGCCGGATCTCACTACTTCCACAGAA ATGGCGGATATTC | Cas1 forward for Gibson assembly pET-His-SUMO/Cas1 |
| 6 | AGATTGGTGGGATGAAAAAGCTATTAAACA CTCTATATGTGACTCAGC | Cas1 reverse for Gibson assembly pET-His-SUMO/Cas1 |
| 7 | TTAATAGCTTTTTCATCCCACCAATCTGTTCT CTGTGAGC | pET-His-SUMO forward for Gibson assembly pET-His-SUMO/Cas1 |
| 8 | ATTTCTGTGGAAGTAGTGAGATCCGGCTGCT AACAAAGCC | pET-His-SUMO reverse for Gibson assembly pET-His-SUMO/Cas1 |
| 9 | AACTTTAAGAAGGAGATATACATATGGCCA GTAATGAAGAAGACCGCTATTTG | Cas4 forward for Gibson assembly pET/Cas4 Cter $His_6$ |
| 10 | GTGATGGTGATGGGAGCCTTCGCTCAGTCTC CCCTCAATG | Cas4 reverse for Gibson assembly pET/Cas4 Cter $His_6$ |
| 11 | AGACTGAGCGAAGGCTCCCATCACCATCAC CATCACTGACTCGAGTTAACCTAGGCTGC | pET forward for Gibson assembly pET/Cas4 Cter $His_6$ |
| 12 | GCGGTCTTCTTCATTACTGGCCATATGTATA TCTCCTTCTTAAAGTTAAAC | pET reverse for Gibson assembly pET/Cas4 Cter $His_6$ |
| 13 | GCGCACTTTCAGTTTTGCAAACGACAG | Forward for Cas4 Q16A |
| 14 | GAGTCCCGACAACATCAAATAGCGG | Reverse for Cas4 Q16A; Revesre for Cas4 H17A |
| 15 | CAGGCGTTTCAGTTTTGCAAACGACAG | Forward for Cas4 H17A |

| | | |
|---|---|---|
| 16 | GCGTGCAAACGACAGTGGGCCTTGATTC | Forward for Cas4 F20A |
| 17 | CTGAAAGTGCTGGAGTCCCGACAA | Reverse for Cas4 F20A |
| 18 | GCGTGGGCCTTGATTCACATCGAGCAGC | Forward for Cas4 Q24A |
| 19 | TCGTTTGCAAAACTGAAAGTGCTGGAG | Reverse for Cas4 Q24A |
| 20 | GCGGAAGAGAATGTCAGGACGATTG | Forward for Cas4 W34A |
| 21 | CTGCTGCTCGATGTGAATCAAGG | Reverse for Cas4 W34A |
| 22 | GCGCATTTGCATAAAAAAGCCGACCAACCG | Forward for Cas4 Q44A |
| 23 | CCCTTCAATCGTCCTGACATTCTC | Reverse for Cas4 Q44A |
| 24 | GCGCTGCAATCTATTTGCTTGCCG | Forward for Cas4 S194A |
| 25 | GCAATTGTTGCAAAAAGGGCCTGTC | Reverse for Cas4 S194A |
| 26 | GCGGGGCAGGCGGCGATCAACTATAATAAAG | Forward for Cas1 E (E166A) |
| 27 | CCATCCACGTAAACTTTCTAAGCTGTCAC | Reverse for Cas1 E (E166A) |
| 28 | TGAAAGCGGAACTCGCGGCGTTAATTGATGAAGAAAAAGACAGTTTGCGC | Forward for Cas2 α2 (T46A, T49A, L53A, T56A and S57A) |
| 29 | ACGACGCTAATTGCGCTGAATCCACAATGCACTCAAA | Reverse for Cas2 α2 (T46A, T49A, L53A, T56A and S57A) |
| 30 | TGACTCGAGTTAACCTAGGC | Forward for Cas4 ΔCth |
| 31 | GCGTTTATTCATCAGCTTCGGC | Reverse for Cas4 ΔCth |
| 31 | GCGGCGTCTGTGGCGGCGTACATTGAGGGGAGACTGAGCG | Forward for Cas4 Q (K206A, R207A, K210A and R211A) |
| 32 | ATTCATCAGCTTCGGCAAGCAAATAGATTGC | Reverse for Cas4 Q (K206A, R207A, K210A and R211A) |

**Table S2. Substrate oligonucleotides used in this study.** PAM and repeat sequences are in bold.

| Name | Sequence (5´ → 3´) Description | Description |
|---|---|---|
| 1 | CGTAGCTGAGGACCACCAGAACTTTTTT**GAA**TTTTTT | PAM/PAM 22 bp duplex strand 1; PAM/NoPAM 22 bp duplex strand 1; PAM/Proc 22 bp duplex strand 1 |
| 2 | GTTCTGGTGGTCCTCAGCTACGTTTTTT**GAA**TTTTTT | PAM/PAM 22 bp duplex strand 2 |
| 3 | GTTCTGGTGGTCCTCAGCTACGTTTTTTTTTTTTTTT | PAM/NoPAM 22 bp duplex strand 2 |
| 4 | GTTCTGGTGGTCCTCAGCTACGTTTTTT | PAM/Proc 22 bp duplex strand 2; Proc/Proc 22 bp duplex strand 2 |
| 5 | CGTAGCTGAGGACCACCAGAACTTTTTT | Proc/Proc 22 bp duplex strand 1 |
| 6 | CGTAGCTGAGGACCACCAGAACAGTTTTT**GAA**TTTTTTT | PAM/PAM 24 bp duplex strand 1 |
| 7 | CTGTTCTGGTGGTCCTCAGCTACGTTTTT**GAA**TTTTTTT | PAM/PAM 24 bp duplex strand 2 |
| 8 | CGTAGCTGAGGACCACCAGTACTTTTTT**GAA**TTTTTT | HSI substrate prespacer strand with PAM; PAM/NoPAM, PAM/Proc strand 1 for comparison of PAM/NoPAM, PAM/Proc and HSI substrate cleavage |
| 9 | TCAAT**ATTTCAATCCACGCACCCATGAAGAGTGCGAC**AGCGAAAATCCCCTAGATTC | HSI substrate bottom strand with 20 bp leader, 32 bp repeat, 5 bp spacer (used with 19, 20 and 21) |
| 10 | GAATCTAGGGGATTTTCGCT | HSI substrate top strand 20 bp leader (used with 18, 20 and 21) |
| 11 | GTACTGGTGGTCCTCAGCTACGTTTTTT**GTCGCACTCTTCATGGGTGCGTGGATTGAAAT**ATTGA | HSI prespacer integrated at leader repeat site (used with 18, 19 and 20) |
| 12 | GTACTGGTGGTCCTCAGCTACGTTTTTTTTTTTTTTT | PAM/NoPAM strand 2 for comparison of PAM/NoPAM, PAM/Proc and HSI substrate cleavage |
| 13 | GTACTGGTGGTCCTCAGCTACGTTTTTT | PAM/Proc strand 2 for comparison of PAM/NoPAM, PAM/Proc and HSI substrate cleavage |
| 14 | CGTAGCTGAGGACCACCAGAACGGGGTT**GAA**TTTTTTTTTT | 4 GC bp at end of 26 bp duplex with 2nt between duplex and PAM strand 1 |

| | | |
|---|---|---|
| 15 | CGTAGCTGAGGACCACCAGAACGGGATT **GAA**TTTTTTTTTT | 3 GC bp at end of 26 bp duplex with 2nt between duplex and PAM strand 1 |
| 16 | CGTAGCTGAGGACCACCAGAACGGAATT **GAA**TTTTTTTTTT | 2 GC bp at end of 26 bp duplex with 2nt between duplex and PAM strand 1 |
| 17 | CGTAGCTGAGGACCACCAGAACGAAATT **GAA**TTTTTTTTTT | 1 GC bp at end of 26 bp duplex with 2nt between duplex and PAM strand 1 |
| 18 | CGTAGCTGAGGACCACCAGAACAAAATT **GAA**TTTTTTTTTT | 0 GC bp at end of 26 bp duplex with 2nt between duplex and PAM strand 1 |
| 19 | CCCCGTTCTGGTGGTCCTCAGCTACGTT | 4 GC bp at end of 26 bp duplex with 2nt between duplex and PAM strand 2 |
| 20 | TCCCGTTCTGGTGGTCCTCAGCTACGTT | 3 GC bp at end of 26 bp duplex with 2nt between duplex and PAM strand 2 |
| 21 | TTCCGTTCTGGTGGTCCTCAGCTACGTT | 2 GC bp at end of 26 bp duplex with 2nt between duplex and PAM strand 2 |
| 22 | TTTCGTTCTGGTGGTCCTCAGCTACGTT | 1 GC bp at end of 26 bp duplex with 2nt between duplex and PAM strand 2 |
| 23 | TTTTGTTCTGGTGGTCCTCAGCTACGTT | 0 GC bp at end of 26 bp duplex with 2nt between duplex and PAM strand 2 |
| 24 | AATGAACGAAAATTCCCTATTTTATCAAA GTGATTTTCTAGAATCTAGGGGATTTTCG CT**GTCGCACTCTTCATGGGTGCGTGGA TTGAAAT**ATTGA | CRISPR 60 bp leader, 32 bp repeat and 5 bp spacer strand 1 |
| 25 | TCAAT**ATTTCAATCCACGCACCCATGA AGAGTGCGAC**AGCGAAAATCCCCTAGA TTCTAGAAAATCACTTTGATAAAATAGG GAATTTTCGTTCATT | CRISPR 60 bp leader, 32 bp repeat and 5 bp spacer strand 2 (Reverse complement of 24) |
